# TH5487 specifically targets NLRP3 in FCAS patients resistant to MCC950

**DOI:** 10.1101/2025.06.05.658156

**Authors:** Angela Lackner, Lemuel Leonidas, Alijah Macapagal, Hannah Lee, Huilin Xu, Yanfei Qiu, Melissa Campos, Julia E. Cabral, Janset Onyuru, Sanika Kulkarni, Lauren V. Albrecht, Hal M. Hoffman, Reginald McNulty

**Affiliations:** Laboratory of Macromolecular Structure, Department of Molecular Biology & Biochemistry, University of California, Irvine; Department of Developmental and Cell Biology, Charlie Dunlop School of Biological Sciences, University of California, Irvine; Division of Pediatric Allergy, Immunology, and Rheumatology, Rady Children’s Hospital of San Diego, University of California, San Diego, San Diego; Department of Pharmaceutical Sciences, University of California, Irvine

## Abstract

The NLRP3 inflammasome plays a central role in innate immunity and is activated in response to mitochondrial dysfunction and oxidized DNA. Here, we demonstrate that repurposed small-molecule inhibitors originally developed for DNA glycosylases, TH5487 and SU0268, potently inhibit NLRP3 activation *ex vivo* in human Peripheral Blood Mononuclear Cells (PBMCs) with IC_50_ of 1.62 µM and 3.24 µM, respectively. We show that these inhibitors prevent mitochondrial localization of NLRP3 and directly block inflammasome assembly. They also reshape the immune landscape decreasing IL-1β, while increasing IFN-β. Structural and biophysical analyses reveal a two-site DNA binding model in which NLRP3 engages oxidized DNA with a KD1 of 0.268 nM and KD2 3.02 nM. Importantly, these inhibitors block IL-1β secretion in L353P Familial Cold Autoinflammatory Syndrome (FCAS) patient PBMCs where MCC950 fails, demonstrating the therapeutic potential for inflammasome-driven diseases. Together, our findings reveal a novel druggable mechanism of inflammasome inhibition through interference with oxidized DNA sensing and localization, offering new opportunities for treatment of chronic inflammatory disorders.

## MAIN

Activation of NLRP3 inflammasome assembly begins when Toll-like receptors (TLRs) recognize pathogen-associated molecular patterns (PAMPs) and promote nuclear localization and activation of NF-κB^1^. The NOD-like receptor pyrin domain-containing protein 3 (NLRP3) acts as a pathogen and toxicant sensor downstream of NF-κB, orchestrating inflammasome activation and the production of IL-1β in response to sterile tissue injury^2,3^. NLRP3 inflammasome activation follows a biphasic process. First, transcriptional priming through LPS interaction with TLR2/4, leads to NF-κB-mediated upregulation of NLRP3 and pro-IL-1β. Reactive oxygen species (ROS) production in the mitochondria results in cytidine/uridine monophosphate kinase 2 (CMPK2) initiating mitochondrial replication^4^. The required unraveling of mitochondrial DNA (mtDNA) for replication exposes the DNA to ROS, resulting in oxidation of mtDNA. Repair of oxidized DNA bases in the mitochondria by human 8-oxoguanine DNA glycosylase 1 (hOGG1)^5^ or elimination of oxidized DNA through mitophagy^6,7^, inhibits inflammasome activation in this half of the biphasic process. In the second half, foreign and cytosolic signals such as bacterial toxins, microcrystalline substances, ATP, silica^8^, asbestos, alum, and hydroxyapatite (HA)^9^, overwhelm initial inflammasome inhibition and sustained ROS production promoting inflammasome subunit assembly and activation. Since most of the agents that initiate inflammasome activation are dissimilar in structure, it suggests that they trigger a common cellular event. One such event is the production of ox-mtDNA^4^. Ox-mtDNA is cleaved to 500-650 bp fragments by flap endonuclease 1 (FEN1) to be exported into the cytoplasm via mitochondrial permeability transition pores (mPTP) and voltage-dependent anion channels (VDAC). Once in the cytoplasm, oxDNA can associate with NLRP3 promoting assembly and activation^10^. NLRP3 inflammasome pro-caspase-1 undergoes autocleavage where mature caspase-1 yields mature IL-1β, IL-18, and gasdermin D. Upon activation, immune molecules, including mt-OxDNA can be released through gasdermin D pores, signaling to other cells via TLR9 and promoting additional TNF-α cytokine release.

NLRP3 has recently been reported to have glycosylase-like activity in that it can cleave oxidized DNA. The protein fold of NLRP3 is similar to hOGG1^11^ and residues involved in nucleophilic attack of oxidized DNA in hOGG1 are conserved in NLRP3. Drugs that influence activity of hOGG1 with oxidized DNA, also bind to NLRP3 and inhibit inflammasome activation^12,13^. This study aims to decipher a mechanism of drug inhibition for hOGG1 repurposed drugs that inhibit inflammasome activation. We illustrate small molecules that bind to NLRP3 prevent inflammasome assembly and demonstrate that they suppress inflammation in hyperactive states present in Familial Cold Autoinflammatory Syndrome (FCAS) patients harboring the L353P mutation who are resistant to MCC950^14^. These drugs limit the response of NLRP3 to mitochondrial oxidized DNA. Moreover, inhibition of NLRP3 response to oxidized DNA promotes alternative cellular response by c-GAS/STING and type I interferon (IFN-β).

## RESULTS

### SU0268 and TH5487 inhibit inflammasome activation in primary human PBMCs

Glycosylases are involved in base excision repair, where they remove oxidized guanine from DNA. The recent discovery that NLRP3 has glycosylase activity and shares active site residues with hOGG1 suggests that NLRP3 may interact with similar small molecules as hOGG1. Indeed, the hOGG1-targeting small molecules SU0268 and TH5487 have been shown to bind to NLRP3 and inhibit inflammasome activation in mouse macrophages^12^ (Fig. 1A). Given the sequence variations between mouse and human NLRP3, which impact disease outcomes differently^15^, we investigated corresponding drug effects in the human monocyte cell line THP-1 (Fig. S1).

**Fig. 1:**
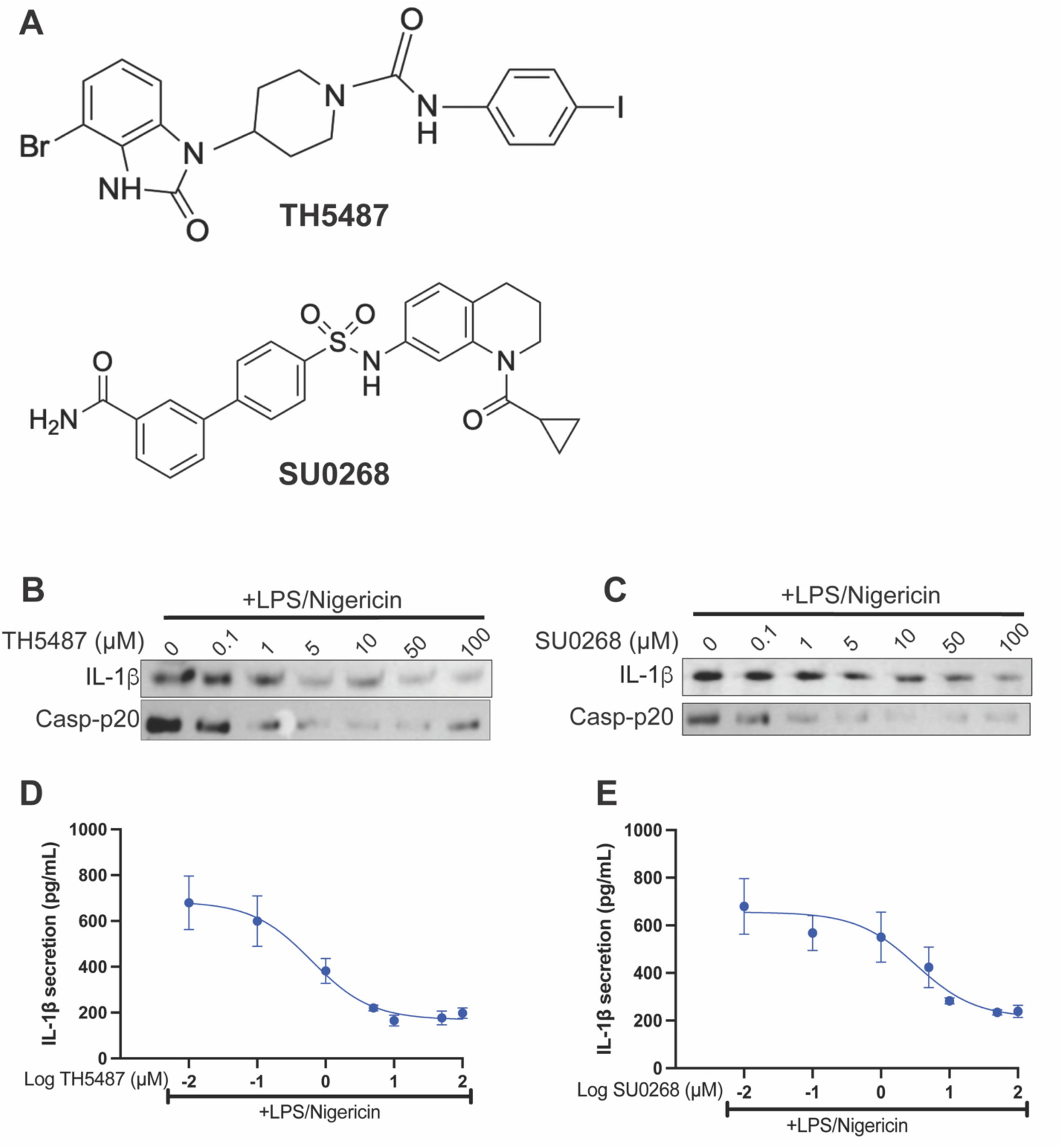
Repurposed OGG1 inhibitors suppress IL-1β and caspase-1 activation in PBMCs from healthy human donors.. A) The chemical structures of two OGG1 inhibitors: TH5487 (top) and SU0268 (bottom) B) Primary human PBMCs from healthy donors were activated with LPS and Nigericin and treated with TH5487 at 0.1-100 µM. IL-1β and Casp-p20 release was visualized by western C) Primary human PBMCs were activated with LPS and Nigericin and treated with SU0268 at 0.1-100 µM. IL-1β and Casp-p20 release was visualized by western blot. D) IL-1β release from TH5487 challenged cells was quantified by ELISA. IC50_TH5487_: 1.62 µM, R^2^_TH5487_: 0.9203. E) IL-1β release from SU0268 challenged cells was quantified by ELISA. IC50_SU0268_: 3.25 µM, R^2^_SU0268_: 0.8209. For both graphs, error bars signify the mean +/- SD. The data was analyzed by non-linear regression.

Since hOGG1 activation promotes the repair of oxidized mitochondrial DNA^5,16,17^, which subsequently inhibits inflammasome activation^5,18^, selective targeting of hOGG1 by these small molecules would theoretically increase ROS, leading to enhanced inflammasome activation. However, because these molecules also bind to NLRP3, we wanted to confirm if TH5487 and SU0268 could inhibit inflammasome activation in the complex model of primary human peripheral blood mononuclear cells (PBMCs). These cells from a healthy donor contained a variety of leukocytes commonly found in circulating blood, and were selected based on their high percentage of CD14+ monocytes (Table S1). Primary PBMCs were primed for 2 hours with 1.6 µg/mL LPS and activated for 1 hour with 20 µM Nigericin, and the supernatant of the cells was probed for the inflammasome-associated cytokine IL-1β as well as the cleaved form of Caspase-1(Fig. 1B-C). We found that TH5487 and SU0268 inhibited IL-1β release. The 1 µM dose of TH5487 inhibited IL-1β secretion by 51.8%, and 5 µM SU0268 inhibited IL-1β and 37.7% %, when quantified by ELISA (Fig. 1D-E). Caspase-1 p-20 secretion was also reduced upon treatment with both inhibitors (Fig.S2 A-B). Cytotoxicity measurements revealed that these respective doses also reduced overall cell toxicity by 12.6% with 1 µM TH5487 and 11% with 5 µM SU0268 (Fig. S3 A-B). We recapitulated this experiment in immortalized human THP-1 monocyte cells, priming them for 16 hours with 400 ng/mL LPS and then activating with 20 µM Nigericin. As expected, both drugs showed dose-dependent NLRP3 inflammasome inhibition and inhibition of IL-1β and caspase-1 p20 secretion with lower doses of TH5487 and SU0268 (Fig. S1, Fig. S2D-E, and Fig. S4A-B). A 0.1 µM dose of TH5487 and a 1 µM dose of SU0268 inhibited IL-1β secretion by 63.2% and 57.3%, respectively (Fig. S1). We confirmed that these inhibitors work similarly with activation by Nigericin or ATP, in immortalized mouse macrophages, and did not show an increase in cytotoxicity, showing robust inhibition across many cell models (Fig.S5 & S6 A-D).

### Small molecule drugs prevent assembly of NLRP3 inflammasome subunits

To further investigate the mechanism of inhibition, we examined the effect of these drugs on inflammasome assembly. We hypothesized that these small molecules would interfere with the formation of the inflammasome complex. We stimulated human monocytes (THP-1s) in the presence of either LPS/Nigericin or LPS/ATP along with increasing concentrations of SU0268 and TH5487. Immunoprecipitation against NLRP3 was then performed to assess assembly of key inflammasome subunits: pro-caspase-1, NEK7, and ASC.

In monocytes, we found 0.1 µM TH5487 was sufficient to reduce the amount of ASC associated with NLRP3 by 77.3% (Fig. 2A,B). The ser/thr kinase NEK7 showed an increase upon addition of ATP alone, but no significant difference in NEK7 association with NLRP3 was achieved with 0.1-100 µM TH5487 (Fig. 2A,C). Pro-caspase-1 decreased by 78.2% with 5 µM TH5487 (Fig. 2D). The same amount of lysate was loaded for each treatment, showing no change in NLRP3 levels (Fig. 2A). As little as 0.1µM SU0268 caused 81.3% reduction in ASC with NLRP3 (Fig. 2E). Unlike TH5487, 10 µM SU0268 decreased NEK7 association by 35.7% (Fig. 2F). SU0268 outperformed TH5487 in preventing association with pro-caspase-1. A 46% reduction in pro-caspase-1 occurred with 0.1 µM SU0268 (Fig. 2G). The observed decrease in NEK7 binding, which can interact with NLRP3 independently of ASC, highlights the specificity of the drugs in affecting proteins that directly bind NLRP3 within the inflammasome complex and related partners. We further validated this data using immunofluorescence staining to show that both 50 and 100 µM of SU0268 efficiently inhibited the association between NLRP3 and ASC compared to control cells, and confirmed that these drugs inhibited ASC speck formation, a hallmark of NLRP3 inflammasome activation (Fig. 2H, S7)^19,20^. Treatment of with 50 µM SU0268 reduced ASC speck formation by 75.8% compared to controls, and 100 µM SU0268 further reduced ASC speck formation by a total of 90.6% (Fig. 2I). We confirmed that this reduction was not simply caused by a decrease in protein expression in response to the drug treatment (Fig. S8A). We confirmed that the expression of NLRP3, ASC, or OGG1 proteins was not affected by treatment of SU0268 from 0.1-100 µM by western blot and RT-qPCR (Fig. S8B-F). These results indicate that the drugs impact NLRP3 interactions with both oxidized DNA^12^ and inflammasome subunits, confirming the link between NLRP3’s oxidized DNA interactions and its ability to form an active inflammasome complex.

**Fig. 2:**
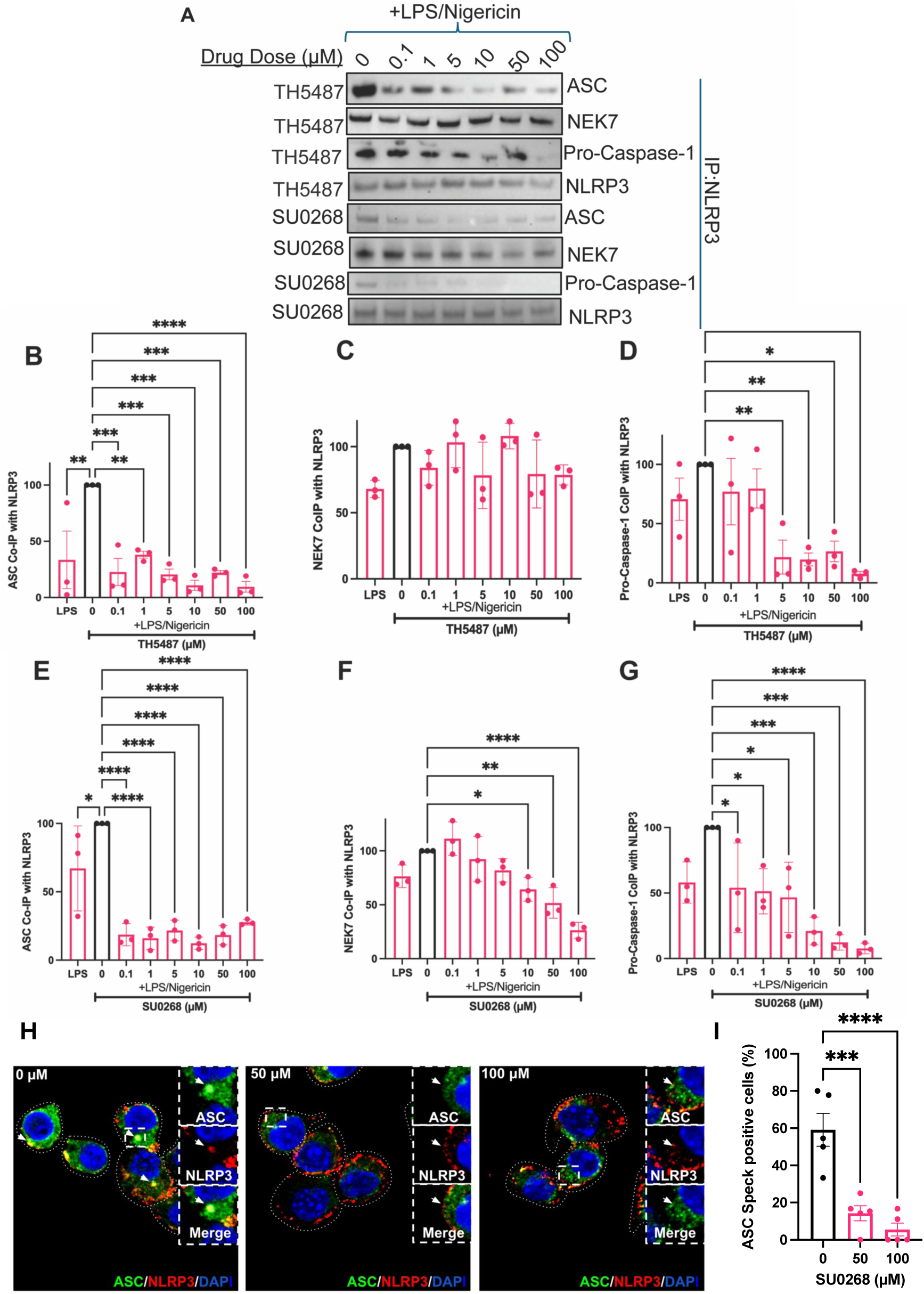
TH5487 and SU0268 disrupt inflammasome assembly by blocking NLRP3-dependent recruitment of ASC, NEK7, and pro-caspase-1. A) THP1 cells activated with LPS and ATP were treated with either TH5487 or SU0268 at 0.1-100 µM. NLRP3 was isolated using protein G magnetic beads and western blots on the fraction that co-immunoprecipitated with NLRP3 were run as western blots. The following graphs represent the quantified amount of ASC B) NEK7 C) and Pro-caspase-1 D) that co-immunoprecipitated with NLRP3 in primed, activated, and inhibited with TH5487. The following graphs represent the quantified amount of ASC E) NEK7 F) and Pro-caspase-1 G) that co-immunoprecipitated with NLRP3 in primed, activated, and inhibited with SU0268. H) ASC speck formation was visualized by immunofluorescence. Immortalized BMDMS were primed with LPS and activated with ATP. ASC was stained green, NLRP3 was stained green, and the nucleus was stained blue. Left: No drug treatment, Middle: 50 µM SU0268 treatment, Right: 100 µM SU0268 treatment I) Quantification of the percentage of ASC speck containing cells from (H) For all graphs, error bars signify the mean +/- SEM. The data was analyzed by one-way ANOVA with N=3. p****<0.0001, p***<0.001, p**<0.01, p*<0.05.

### Dual Inhibition of NLRP3 Priming and TNF-α Release by TH5487 and SU0268

Since TH5487 and SU0268 also directly target hOGG1^21,22^, we hypothesized that they would inhibit the priming step of inflammasome activation, as hOGG1 can increase NF-κB activity^23,24^, driving the production of priming cytokines like TNF-α^24^. Indeed, we found that TNF-α expression, elevated with LPS/ATP treatment, was reduced in the presence of TH5487 or SU0268 (Fig. S9A-B). A concentration of 5 µM TH5487 reduced TNF-α expression by 21% (Fig. S9A). Similarly, 5 µM SU0268 reduced TNF-α expression by 35% compared to LPS/ATP treatments (Fig. S9B). These results demonstrate that these small molecules effectively target both the priming and activation phases of inflammasome activation.

We also wanted to explore if TNF-α signaling was also reduced in these conditions. To do this, the supernatant of LPS/ATP-activated cells challenged with TH5487 or SU0268 was probed using a western blot for TNF-α secretion (Fig. S9C). A concentration of 0.1 µM TH5487 reduced TNF-α release by 35.2%. (Fig. S9D). Similarly, 0.1 µM SU0268 reduced TNF-α release by 25.9% compared to LPS/ATP treatments (Fig. S9E). A higher dose of 50 µM TH5487, reduced TNF-α secretion further by 55.6%. However, a more sensitive response was achieved with 5 µM SU0268 which caused TNF-α reduction by 45.5%. Both drugs inhibit TNF-α production and signaling, but lower concentrations of SU0268 bring about higher inhibitory effects. Our data supports previous studies showing TH5487 and SU0268 inhibit LPS-induced TNF-α, due to their dual role in inhibiting hOGG1^21,22^.

### TH5487 and SU0268 inhibit NLRP3 activation and mitochondrial association induced by mitochondrial dysfunction

Antimycin A (AA) inhibits mitochondrial complex III, leading to electron transport chain disruption and increased production of mitochondrial ROS^25^. Mitochondrial membrane potential perturbations are known to promote the release of oxidized mitochondrial DNA (ox-mtDNA) and induce NLRP3 inflammasome activation. Prior studies, including those by the Subramanian group, have shown that NLRP3 associates with mitochondria via mitochondrial antiviral signaling protein (MAVS)^26^. To test if NLRP3 activation driven by mitochondrial stress could be inhibited by repurposed glycosylase small molecules, we treated immortalized THP-1 cells with 400 ng/mL LPS for 16 hours followed by stimulation with 10 µM AA for 2 hours. AA robustly induced IL-1β secretion, which was suppressed by TH5487 and SU0268 (Fig. 3A). Specifically, 5 µM TH5487 and 0.1 µM SU0268 reduced IL-1β secretion by 68.2% and 52.8%, respectively (Fig. 3B-C). Furthermore, AA promoted NLRP3 localization to mitochondria, and this mitochondrial translocation was markedly reduced by both inhibitors (Fig. 3D). As little as 5 µM TH5487 reduced NLRP3 mitochondrial localization by 63.9%, and 0.1 µM SU0268 reduced NLRP3 mitochondrial localization by 70% (Fig. 3E-F). Importantly, total cytosolic NLRP3 levels were not significantly altered by these treatments (Fig. S10), suggesting that a small, regulated pool of NLRP3 is targeted to mitochondria upon activation. These findings support a model in which ox-mt DNA facilitates NLRP3 recruitment to mitochondria, and this process is disrupted by glycosylase inhibitors targeting NLRP3-oxDNA interactions.

**Fig. 3:**
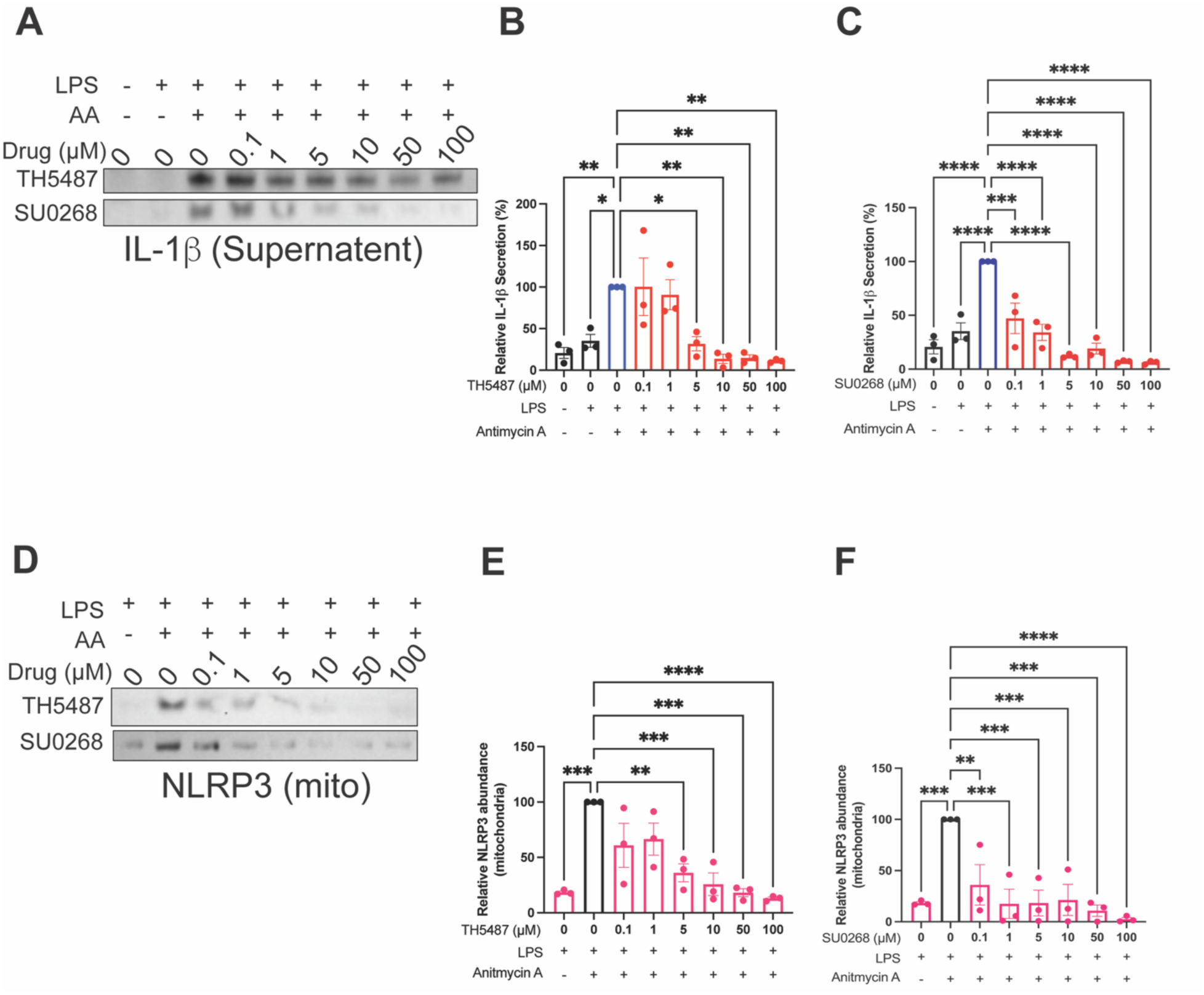
Repurposed inhibitors block IL-1β release and NLRP3 mitochondrial localization under mitochondrial stress. A) THP1 cells activated with LPS and Antimycin A were treated with either TH5487 or SU0268 at 0.1-100 µM. Relative IL-1β release was visualized by western blot. B) Quantification of (A) for TH5487. C) Quantification of (A) for SU0268. D) THP1 cells activated with LPS and Antimycin A were treated with either TH5487 or SU0268 at 0.1-100 µM. The mitochondrial fraction was isolated and the relative amount of NLRP3 associated with the mitochondriavwas visualized by western blot. E) Quantification of (D) for TH5487. F) Quantification of (D) for SU0268.

### Inhibiting NLRP3 interaction with Ox-mtDNA promotes compensation by c-GAS/STING

Stimulation of monocytes with LPS/ATP or LPS/nigericin leads to alterations in the membrane potential of the outer membrane and within intracellular organelles, including the ER, Golgi, and mitochondria^27,28^. These hyperpolarization changes at the outer membrane induce ionic fluxes in intracellular organelles, resulting in abnormal fluctuations in sodium, potassium, calcium, and other signaling molecules^29–31^. Although TH5487 and SU0268 have been shown to decrease the release of oxidized mitochondrial DNA (Ox-mtDNA)^12^, complete inhibition of NLRP3 leads to a cytosolic increase of DNA that is not mitochondrial in origin^12^.

We hypothesize that these significant ionic fluxes within the cell could eventually lead to disturbances and the secretion of nuclear DNA, which may become oxidized and released into the cytosol. Thus, inhibition of NLRP3 is expected to increase cytosolic DNA levels, specifically from nuclear, not mitochondrial, sources. Exposure of oxidized and non-oxidized nuclear DNA in the cytosol could subsequently activate the cGAS/STING pathway^32,33^. We found that NLRP3^-/-^ have decreased oxidized DNA in the mitochondrial compartment compared to wildtype. However, there was no significant difference in oxidized DNA released to the cytosol for NLRP3^-/-^ compared to wildtype (Fig. 4A). This suggests that, in response to a loss of NLRP3, the amount of cytosolic oxidized DNA may be compensated by DNA from that is of nuclear origin. We validated this by measuring nuclear DNA content in the cytosol of THP-1 cells stimulated with LPS and Nigericin and treated with either TH5487 or SU0268. Both inhibitors significantly increased the relative amount of cytosolic nuclear DNA by 5.6-fold or 7.8-fold with 1 µM of TH5487 or SU0268, respectively (Fig. 4B). We then checked for c-GAS/STING to compensate for the increase in oxidized DNA when NLRP3 is inhibited in the presence of LPS/Nigericin. Indeed, we observed a concentration-dependent increase in phosphorylated STING (p-STING) with escalating levels of TH5487, confirming that NLRP3 inhibition triggers cGAS/STING activation (Fig. 4C). The amount of p-STING increased by 43% with 1 µM TH5487 (Fig. 4D), while 5 µM of SU0268 increased the amount of p-STING by 93% (Fig. 4E). Activation of c-GAS/STING is known to lead to type 1 interferon response^1,33,34^. We found 0.1 µM TH5487 caused 107.5% increase in cytosolic IFN-β when stimulated with LPS/nigericin (Fig. 4F). Less sensitivity was seen with 50 µM SU0268, which caused 101.9% increase in IFN-β (Fig. 4G). To determine whether these findings extend to a more physiological model, we repeated experiments using primary human peripheral blood mononuclear cells (PBMCs). PBMCs were primed for 2 hours with 1.6 µg/mL LPS and activated for 1 hour 20 µM Nigericin and challenged with either TH5487 or SU0268. We first evaluated IFN-β expression by RT-qPCR across a range of drug concentrations (0.1-100 µM). TH5487 and SU0268 increased IFN-β transcript levels by 10.7% (5 µM) and 7% (10 µM), respectively (Fig. 4H, S11A&C). To assess whether this transcriptional upregulation translated to IFN-β secretion, we performed ELISA on the supernatant. Treatment with 50 µM TH5487 and SU0268 increased IFN-β secretion by 1.6-fold and 3.7-fold, respectively (Fig. 4I, S11B&D), confirming that inhibition of NLRP3 with these repurposed inhibitors promotes type I interferon responses through the cGAS/STING axis.

**Fig. 4:**
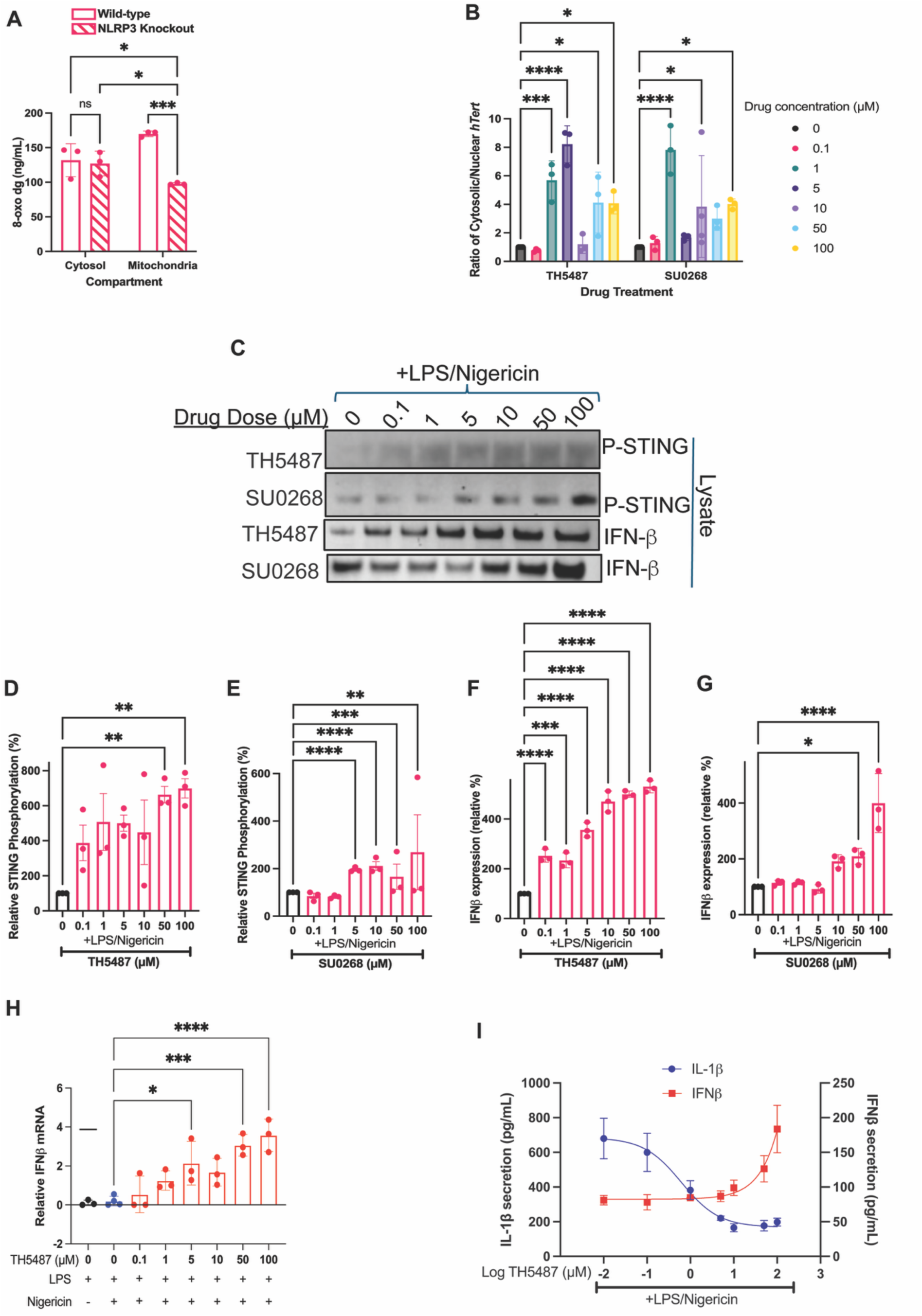
NLRP3 inhibition promotes cGAS-STING activation and IFN-β production in primary human PBMCs. A) NLRP3 wild type and knockout cells were treated with LPS and ATP, the mitochondrial and cytosolic fractions were isolated, and the amount of oxidized DNA in each compartment was quantified by and 8-oxo-dG ELISA. B) THP1 cells activated with LPS and Nigericin were treated with either TH5487 or SU0268 at 0.1-100 µM. The cytosolic and nuclear fraction of DNA were isolated and qPCR was run against *hTert* as a nuclear probe. The ratio of *hTert* in the cytosol to *hTert* in the nucleus was quantified for each drug treatment. C) THP1 cells activated with LPS and Nigericin were treated with either TH5487 or SU0268 at 0.1-100 µM. Relative amounts of IFN-β and phosphorylated STING in the lysate were analyzed by western blot. The relative amount of secreted IFN-β (D/E) or internal phosphorylated sting (F/G) was quantified for both drugs. H) Primary human PBMCs were activated with LPS and Nigericin and treated with TH5487 at 0.1-100 µM and IFN-β expression was quantified with qPCR. I) Primary human PBMCs were activated with LPS and Nigericin and treated with TH5487 at 0.1-100 µM. IFN-β and IL-1β release was quantified by ELISA. IC50_TH5487_: 1.62 µM, R^2^_TH5487_: 0.9203, EC50_TH5487_: 53.2 µM, R^2^_TH5487_: 0.9203. For bar graphs, error bars signify the mean +/- SEM and the data was analyzed by one-way ANOVA with N=3, where p****<0.0001, p***<0.001, p**<0.01, p*<0.05. For the line graph, error bars signify mean +/- SD and the data was analyzed using non-linear regression.

### Biophysical analysis reveals NLRP3 binds DNA in a two-site binding model

Given our physiological data supporting the therapeutic relevance of inhibiting NLRP3:DNA interactions, we sought to further define this relationship biophysically. We adapted a method from the Nogales group using streptavidin-coated affinity cryo-electron microscopy (cryo-EM) grids^35^. Full-length NLRP3 was first incubated with biotinylated, non-oxidized mitochondrial DNA (bt-DNA) to form a stable NLRP3–bt-DNA complex, which was then applied to the streptavidin-coated grid surface (Fig. S12A). We used non-oxidized DNA because it binds NLRP3 but is not cleaved, as previously reported. NLRP3 was purified as a decamer in complex with TH5487 and confirmed as a single high-order species by SDS-PAGE, western blot, and native gel electrophoresis (Fig. S12B). After applying the NLRP3–bt-DNA complex to the affinity grids, particles were visualized using a Falcon 4i microscope. While particles of various sizes were present, the underlying streptavidin lattice was visible in the background. Since the Fourier transform of a lattice is another lattice, the reciprocal lattice spots were visible in the FFT (Fig. S12C-D). Computational subtraction of the streptavidin lattice markedly improved particle visualization and eliminated FFT contamination^36^ (Fig. 5A, S12E). This revealed two main populations of particles: a smaller class around ∼100 Å and a larger class between 180–200 Å, consistent with the published cryo-EM structure of NLRP3 bound to the small-molecule inhibitor MCC950.

**Fig. 5:**
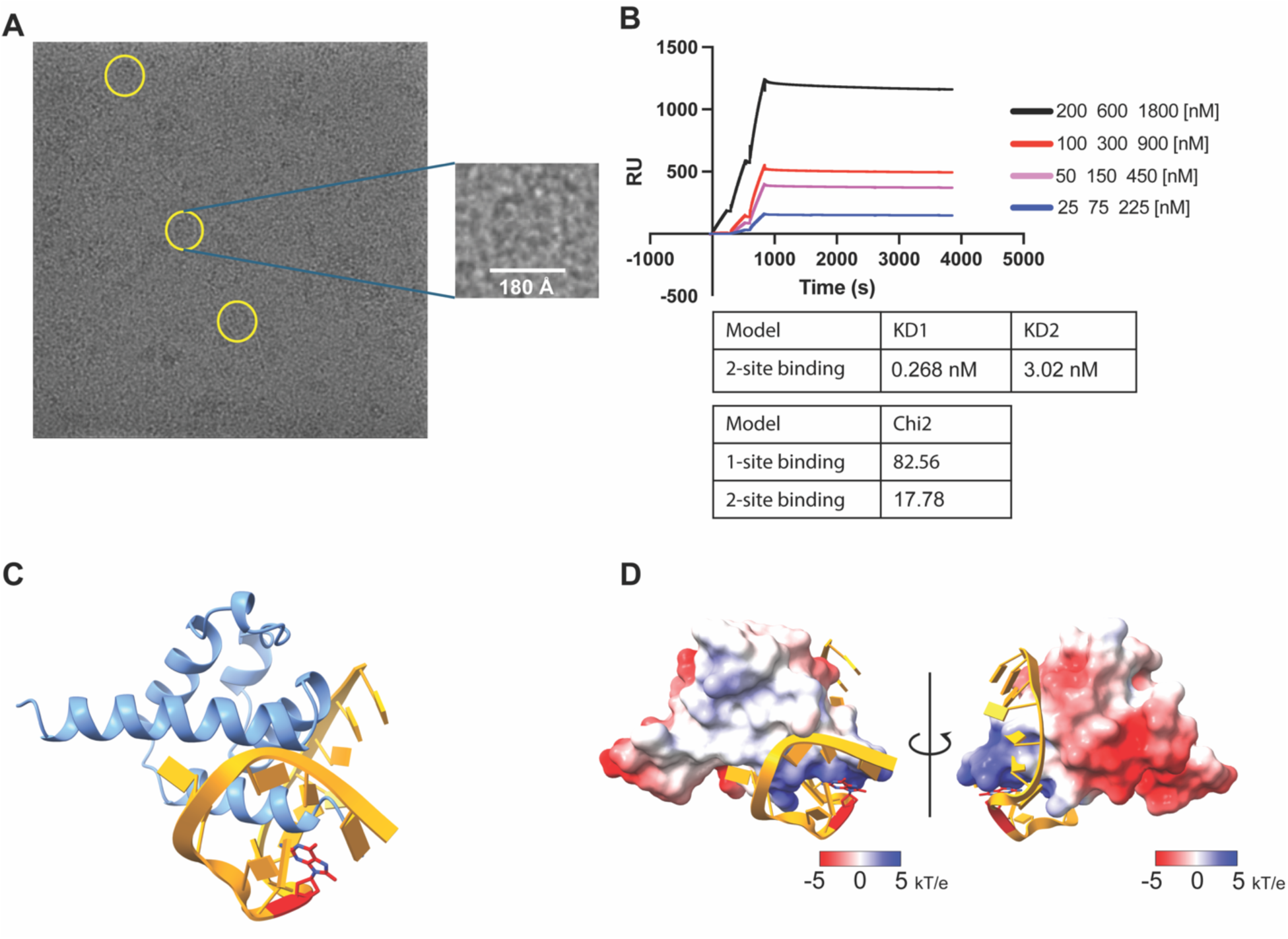
NLRP3 binds oxidized mitochondrial DNA through a two-site mechanism revealed by cryo-EM and SPR. A) Example cryoEM micrograph after streptavidin lattice removal. Yellow circles show NLRP3 particles bound to the streptavidin/biot-DNA layer. Particles measure ∼18nm in diameter, similar to the size of the published inactive decamer of NLRP3 (PDB 7PZC). B) SPR sensogram for quantitative analyses of FL NLRP3 binding 20 base-pair biot-ox-mtDNA. Best fit 2-site-binding model KD values and Chi squared values for both a 1-site and 2-site binding model. C) An AlphaFold model of the NLRP3 pyrin domain (residues 1-85, blue) interacting with single stranded (orange) oxidized DNA (red, from PDBID: 1EBM). D) The electrostatic potential of the NLRP3 pyrin domain AlphaFold model shows a more negative charge in the region closest to the oxidized base (red) in the single stranded DNA (orange). The model has been rotated horizontally 90 degrees to show two different views.

To further characterize the interaction between NLRP3 and DNA, we conducted surface plasmon resonance (SPR) utilizing full-length NLRP3 as an analyte and biot-ox-mtDNA as a conjugated ligand. Our data revealed that NLRP3 bound ox-mtDNA in a dose-dependent manner, and that the data fit much better into a 2-site binding model than into a 1-site binding model, with KD1 and KD2 of 0.268 nM and 3.02 nM (Fig. 5B). Similar to OGG1 which can weakly interact with DNA until it locks on an 8-oxo-DG, we propose the NACHT domain serves to bind DNA weakly (KD2) and the pyrin domain has a higher affinity for 8-oxodDG-containing-DNA (KD1). Using AlphaFold, we computationally confirmed this data, showing that single-stranded (ss) oxDNA, but not non-oxidized ssDNA, can bind the pyrin domain (Fig. 5C-D, S13).

### hOGG1 activator TH10785 binds NLRP3 and inhibits inflammasome activation in primary human PBMCs

The drugs TH5487 and SU0268 inhibit hOGG1, and we demonstrate herein that they also inhibit NLRP3. From a therapeutic perspective, inhibiting hOGG1 could be beneficial in allowing an increase in cytosolic oxidized DNA, which may enhance type-1 interferon immune response. Conversely, if reducing an overactive immune response is desirable, promoting DNA repair by activating hOGG1 while simultaneously inhibiting NLRP3 inflammasome activation could be advantageous. Given that TH5487 and SU0268 inhibit hOGG1, we sought a drug capable of activating hOGG1. TH10785 acts as an hOGG1 activator, initiating DNA repair to mitigate DNA damage and reduce ROS production^37^ (Fig. 6A). So, we investigated if TH10785 could bind NLRP3 and inhibit inflammasome assembly. We found superposition of NLRP3 pyrin and hOGG1 bound to TH10785 showed a similar binding pocket for TH10785 (Fig. 6B). The model was further improved using SWISS MODEL (Fig. 6C). The model was further improved using SWISS MODEL (Fig. 6C). Many homologous residues for the catalytic activity in hOGG1 moved into closer proximity with one another, namely the Lys2 in NLRP3 corresponding to hOGG1’s catalytic Lys249 moved to only 0.632 Å away, the NLRP3’s Asp21 moved 1 Å closer to hOGG1’s charge-donating Asp268, and NLRP3’s Phe75 moved to less than 0.7 Å away from hOGG1’s stabilizing Phe319 (Table S2). Next, we tested whether TH10785 could inhibit NLRP3 inflammasome activation in primary human PBMCs primed and activated with LPS/Nigericin and challenged with 0.1-100 µM of TH10785. We probed the supernatant for IL-1β and Caspase-p20 release (Fig. 6D). As little as 5 µM of TH10785 inhibited NLRP3-dependent IL-1β by 41.4% when quantified by ELISA (Fig. 6E). Caspase-p20 secretion was also reduced (Fig. S2C). We conducted the same experiments to evaluate cGAS-STING pathway activation in cells inhibited with TH10785. We saw a 9.5-fold increase in IFN-β expression by RT-qPCR with 10 µM TH10785, and a 3.8-fold increase in IFN-β secretion by ELISA with 100 µM TH10785 (Fig. 6E-F). This supports cGAS-STING inflammatory compensation in response to NLRP3 inhibition, which may be partially due to increased cytotoxicity of the drug at high doses (Fig. S3C).

**Fig. 6:**
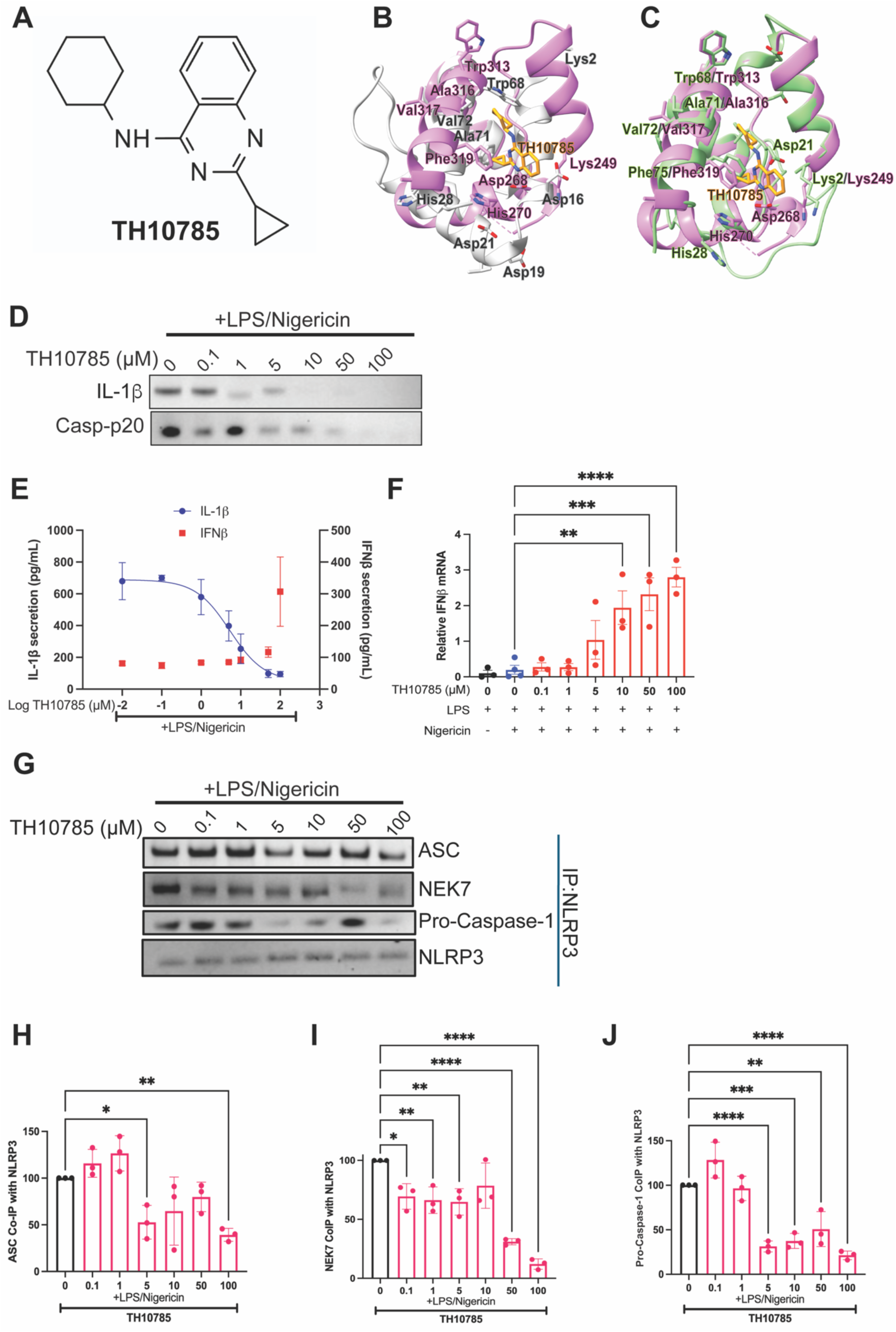
TH10785, a known hOGG1 activator, unexpectedly inhibits NLRP3 and promotes interferon signaling. A) The chemical structure of the OGG1 activator TH10785. B) The NLRP3 pyrin domain (grey, PDBID: 7PZC) docked into the structure of hOGG1 bound to TH10785 (pink, PDBID: 7AYY). C) SWISS-MODEL projection of NLRP3 pyrin based on hOGG1 bound to Ox-DNA (green) docked into the structure of hOGG1 bound to TH10785 (pink, PDBID: 7AYY). D) Primary human PBMCs were activated with LPS and Nigericin and treated with TH10785 at 0.1-100 µM. IL-1β and Casp-p20 release was visualized by western. E) IFN-β and IL-1β release were quantified by ELISA. IC50_TH10785_: 5.329 µM, R^2^_TH10785_: 0.9031. F) IFN-β expression was quantified with qPCR. G) THP1 cells activated with LPS and ATP were treated with either TH10785 at 0.1-100 µM. NLRP3 was isolated using protein G magnetic beads and western blots on the fraction that co-immunoprecipitated with NLRP3 were run as western blots. The following graphs represent the quantified amount of ASC (H) NEK7 (I) and Pro-caspase-1 (J). For bar graphs, error bars signify the mean +/- SEM, and the data was analyzed by one-way ANOVA with N=3, where p****<0.0001, p***<0.001, p**<0.01, p*<0.05. For the line graph, error bars signify mean +/- SD and the data was analyzed using non-linear regression.

### TH10785 binds NLRP3 and inhibits inflammasome activation and assembly in monocytes

To test if inflammasome activity could be inhibited by TH10785 in human monocytes, THP-1 monocytes were primed and activated with LPS/nigericin and treated with 0.001-100 µM TH10785. We evaluated overall inflammasome activity both intercellularly and by IL-1β and Casp-p20 secretion (Fig. S14 A-B, S2F, S4C). As little as 0.1 µM of TH10785 inhibited NLRP3-dependent IL-1β by approximately 50% (Fig. S14B). Similar results were seen in mouse iBMDM cells activated with LPS and ATP and treated with TH10785 (Fig. S14 C-F). We also wanted to examine the mechanism of TH10785-mediated IL-1β decrease. We performed co-IP of NLRP3 with ASC, procaspase-1, and NEK7 (Fig. 6G). We found 5 µM TH10785 decreased the association of ASC, procaspase-1, and NEK7 by 47.7%, 38.2%, and 67.4%, respectively (Fig. 6H-J). These results are significant because they illustrate that we can promote or decrease priming, while simultaneously inhibiting inflammasome activation.

### TH5487 inhibits IL-1β secretion from disease-associated L353P CAPS human PBMCs

Given the need for effective NLRP3 inhibitors in autoinflammatory diseases, we tested the ability of TH5487 to inhibit NLRP3-dependent IL-1β secretion in primary human PBMCs isolated from patients harboring the L353P CAPS mutation. This mutation causes the familial cold and flu syndrome (FCAS) subtype of CAPS, characterized by cold-induced inflammasome hyperactivation and IL-1β overproduction. We sought to compare our results to the well-characterized NLRP3 inhibitor MCC950, which is ineffective at blocking NLRP3 activity in CAPS patients (Fig. 7A)^14^. Primary human PBMCs from L353P patients were primed for 3 hours with 1.6 µg/mL LPS and treated with TH5487 at 0.1-100 µM for 1 hour, and IL-1β release was visualized by western blot (Fig. 7B). Since CAPS mutants can harbor an inflammatory response without a secondary activating signal, no ATP or nigericin was added to this study. To compare TH5487 efficacy to MCC950, wild-type and CAPS PBMCs were primed and activated as previously described, treated with 10 µM TH5487 or MCC950, and IL-1β release was visualized by western blot (Fig. 7C). IL-1β secretion was quantified by ELISA, which showed a 61.4% decrease in NLRP3-dependent IL-1β secretion upon treatment with 10 µM TH5487 (Fig. 7D). In these cells, 10 µM MCC950 proved ineffective, but in wild-type PBMCs, 10 µM MCC950 and 10 µM TH5487 both inhibited NLRP3-dependent IL-1β release by 82.5% and 48.6%, respectively (Fig. 7E). These findings highlight the therapeutic potential of this novel inhibitor to overcome MCC950 resistance in CAPS, offering a promising strategy for targeting NLRP3-driven inflammation where current approaches fail.

**Fig. 7:**
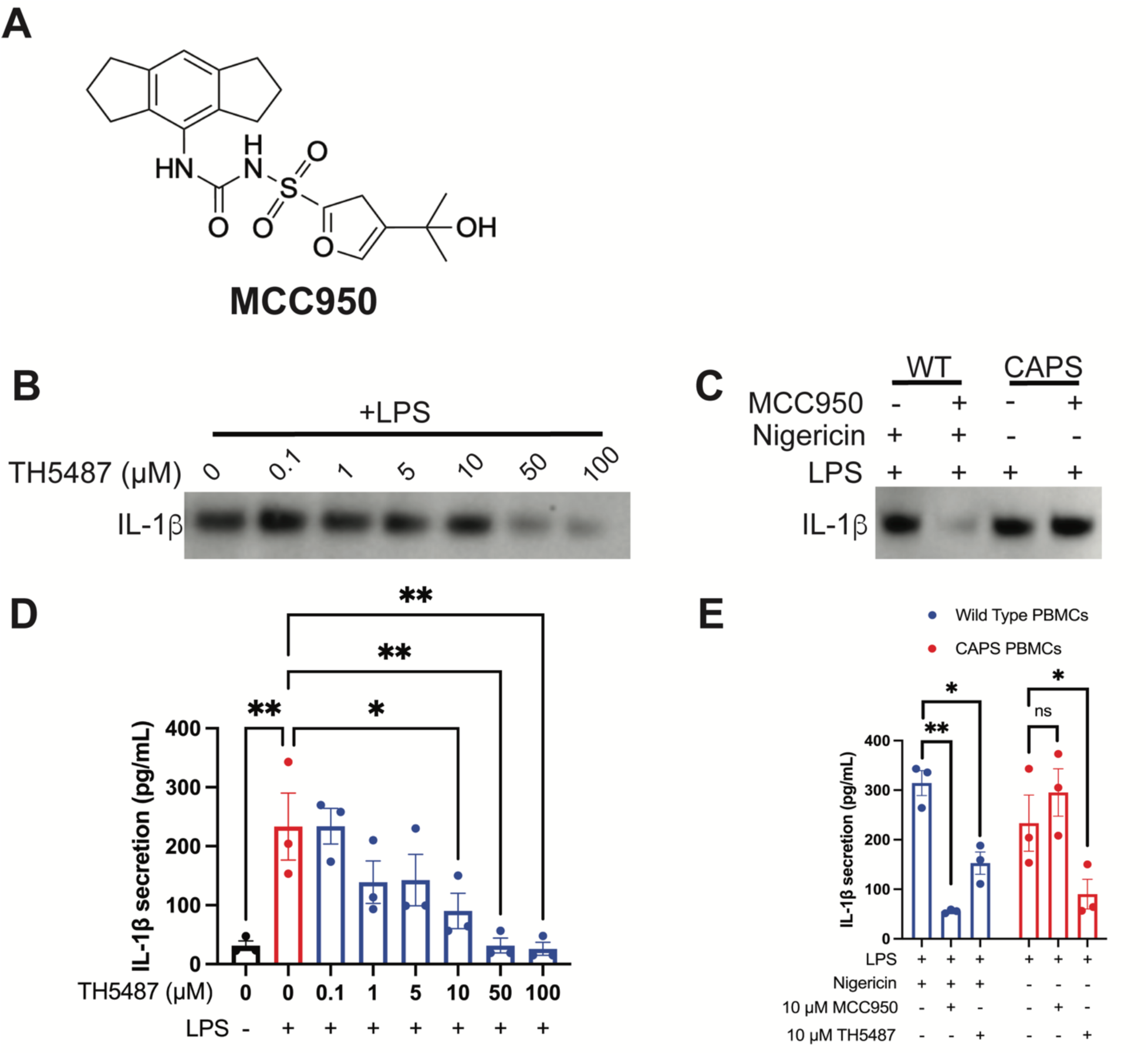
Repurposed inhibitors block inflammasome activation in CAPS PBMCs resistant to MCC950. A) The chemical structure of the NLRP3 inhibitor MCC950. B) Primary human PBMCs from patients harboring the L353P CAPS mutation were activated with LPS and treated with TH5487 at 0.1-100 µM. IL-1β release was visualized by western. C) Primary human PBMCs from healthy donors and patients harboring the L353P CAPS mutation were activated with LPS (CAPS) or LPS and Nigericin (helathy) and treated with TH5487 or MCC950 at 10 µM. IL-1β release was visualized by western. D) IL-1β release from the L353P cells challenged with the dose titration of TH5487 was quantified by ELISA. E) IL-1β release from the healthy or L353P cells challenged with 10 µM TH5487 or MCC950 was quantified by ELISA. For both graphs, error bars signify the mean +/- SEM, and the data was analyzed by one-way ANOVA with N=3, where p**<0.01and p*<0.05.

## DISCUSSION

In summary, this study reveals that drugs that bind human glycosylase hOGG1 can be repurposed to target NLRP3 inflammasome activation in physiologically relevant peripheral blood mononuclear cells (PBMCs) isolated from human blood. Moreover, FCAS patients with L353P that do not respond to MCC950 respond to TH5487. These repurposed inhibitors specifically bind NLRP3 and inhibits inflammasome activation in the presence of antimycin A. If the response was not NLRP3-dependent, one would expect an increase in inflammasome activation when ROS and oxDNA is generated when AA inhibits the mitochondrial electron transport chain. The release of oxidized DNA from the mitochondria and detection in the cytosol is NLRP3 dependent^12^. Herein, we show *NLRP3^-/^*^-^ or NLRP3-drug inhibited states which block release of mitochondrial DNA, can still lead to oxidation and release of DNA that is of nuclear origin. Oxidation of either mitochondrial or nuclear DNA released to the cytoplasm can act as a DAMP. Both forms of oxidized DNA that elicit an inflammation response in this way are linked to aging and age-related pathologies, so-called “inflammaging”^38,39^. We illustrate that inhibition of NLRP3 leads to compensation by c-GAS/STING pathway to induce a type 1 interferon response with production of IFN-β. We propose a general causal mechanism in that inhibiting OGG1-mediated DNA repair causes an increase in cytosolic oxDNA. This increase in cytosolic oxDNA would normally activate NLRP3 inflammasome, but TH5487 and SU0268 prevent NLRP3 from interacting with oxDNA and inflammasome assembly. As we are inhibiting both nuclear and mitochondrial OGG1, an initial cytosolic increase in oxDNA could arise from either source^10^. Our results show an increase in cytosolic oxDNA, which has been shown herein to activate c-GAS/STING and increase IFN-β^40^. We observe a decrease in cytosolic D-loop mtDNA. The total mtDNA decrease suggests mitophagy has occurred which is known to be associated with c-GAS activation^41^. In this case, an initial efflux of ox-mtDNA would activate cGAS/STING and cause mitophagy.

The interplay between IFN-β and IL-1β is complex, as these pathways frequently cross-regulate one another in an antagonistic manner. IFN-β can inhibit inflammasome activity and subsequently reduce IL-1β production by activating signaling cascades that counter inflammasome activation^21,42^. Conversely, IL-1β can suppress IFN-β responses under certain inflammatory conditions, creating an environment that dampens type I interferon signaling^43^.

Chronic inflammation involving IL-1β and the NLRP3 inflammasome has been implicated in various age-related diseases, including Alzheimer’s disease (AD)^44–47^. Although our study does not focus on AD directly, AD serves as a well-studied example of how persistent inflammation, or "inflammaging," can contribute to disease progression. In AD, NLRP3 inflammasome activation occurs in response to amyloid-beta (Aβ) plaques, leading to the release of IL-1β and other pro-inflammatory cytokines that exacerbate neuronal damage^31,45^. Elevated IL-1β levels correlate with synaptic dysfunction and neuronal loss, which are central to cognitive decline^31^. Chronic, low-grade systemic inflammation primes microglia, rendering them more sensitive to Aβ deposits, which could accelerate AD onset and progression in predisposed individuals^48^. Prolonged inflammation, associated with cellular senescence, mitochondrial dysfunction, and oxidative stress, contributes to the premature aging that characterizes inflammaging^49,50^. Consequently, individuals with high levels of chronic inflammation may exhibit physical and cognitive signs of aging earlier than expected.

Studies using NLRP3 knockout models in animals indicate that NLRP3 deficiency reduces neuroinflammation and delays cognitive decline in AD models, underscoring NLRP3’s role in AD pathogenesis^51^. Elevated IL-1β levels in post-mortem AD brain tissues, along with correlations between cerebrospinal IL-1β and cognitive decline, further highlight the relevance of this pathway in human AD^52,53^. Additionally, interactions between Aβ and NLRP3 in microglia create a pro-inflammatory feedback loop, perpetuating neuronal damage and exacerbating disease progression^54^. These insights suggest that targeting the NLRP3 inflammasome to reduce IL-1β release and neuroinflammation could potentially delay the onset or progression of AD and other age-related diseases associated with chronic inflammation^46–48,55^. Selective targeting of the NF-kb induced priming and NLRP3 activation my play critical roles in treatment of inflammaging.

But strictly inhibiting the IL-1β is not the only remedy since production since loss of large abnormal ion concentrations result during pathogen or toxicant and tissue damage can lead to destabilization of mitochondrial and nuclear DNA sources. The ability to inhibit or promote the priming aspect of inflammasome activation while simultaneously inhibiting or promoting repair can have a big impact on inflammaging. The logic is as follows. During mtDNA replication or repair G-rich regions are susceptible to oxidation and DNA strand breaks which can cause deletions^56^. The mitochondrial theory of aging suggest that insufficient availability of peptide synthesis will decrease electron transfer across ETC respiratory complexes ^57^which stall leading to early transfer of electrons to O_2_^58^. The cycle of deletions ^59^ and mutations ^60^ which are uncommon before the age of 40 exponentially accumulate during old age. The accumulation of oxidative damage to mtDNA and nuclear DNA is an important mechanism of aging^61,62^. The mitochondria can control cell fate via efflux of DNA from mPTP (mitochondrial Permeability Transition Pores) to the cytosol^10,63,64^. We find herein the release of mtDNA from the mitochondria is NLR3 dependent. Inhibiting this process stops production of IL-1β, but leads to an increase from nuclear sources. The NLRP3 inhibitor herein, TH10785 which simultaneously activates hOGG1, may facilitate repair of nuclear DNA, leading to reduction in TLR9 induced responses due to the presence of cytosolic nuclear DNA. Given attempt to repair during high ROS can lead to deletions and mutations,

Moreover, alcohol abuse can also have effects. A single dose of ethanol causes extensive degradation and depletion of hepatic mtDNA within 2 hours in mice^65,66^. After 4 days of alcohol admin numerous deletions of mtDNA prevent mitochondrial replication because there are not enough intact copies of templates. This leads to mtDNA depletion for several days until mtDNA is restored once alcohol intake stops in white blood cells which have a quick turnover^67^. Excessive alcohol and fructose have a profound impact on liver mitochondrial dysfunction and NLRP3 expression, which ultimately progress to metabolic dysfunction-associated steatohepatitis (MASH) and hepatocellular carcinoma (HCC)^68^. Future experiments will evaluate the contribution of NLRP3 to these diseases using small molecule inhibitors illustrated herein.

The ability to simultaneously target the priming and activation phases of inflammasome activation represents a therapeutic advantage. Future experiments will involve optimizing the specificity for biphasic targeting as well as defining molecular determinants of NLRP3 recognition for oxidized DNA and drugs described herein^69,70^.

## DATA AVAILABILITY

We will be willing to distribute materials and protocols to qualified researchers in a timely manner.

### Lead contact

Further information and requests for resources and reagents should be directed to and will be fulfilled by the lead contact, Reginald McNulty.

### Materials availability

There were no unique materials generated by this study. Sources for all materials used to generate the data in this manuscript have been noted in the methods.

### Data and code availability

All data are available in the main text or the supplementary materials.

### Study Approval

Studies performed using PBMCs obtained from patients with CAPS received the approval of the University of California Human Research Protection Program committee, and informed consent was obtained from the subjects before the study.

## ACKNOWLEDGMENTS

Funding: This work was supported by the National Institutes of Health NIAID grant K22AI139444 to R.M.

## AUTHOR CONTRIBUTIONS

Conceptualization: AL, HH, RM

Methodology: AL, RM, MC, LA

Investigation: AL, JC, YQ, LL, AM, HL, HX, SK, MC

Analysis: AL, YQ, LL, HL, MC, LA

Supervision: RM

Writing—original draft: AL, RM

Writing—review & editing: AL, RM

---

**Fig. S1:**
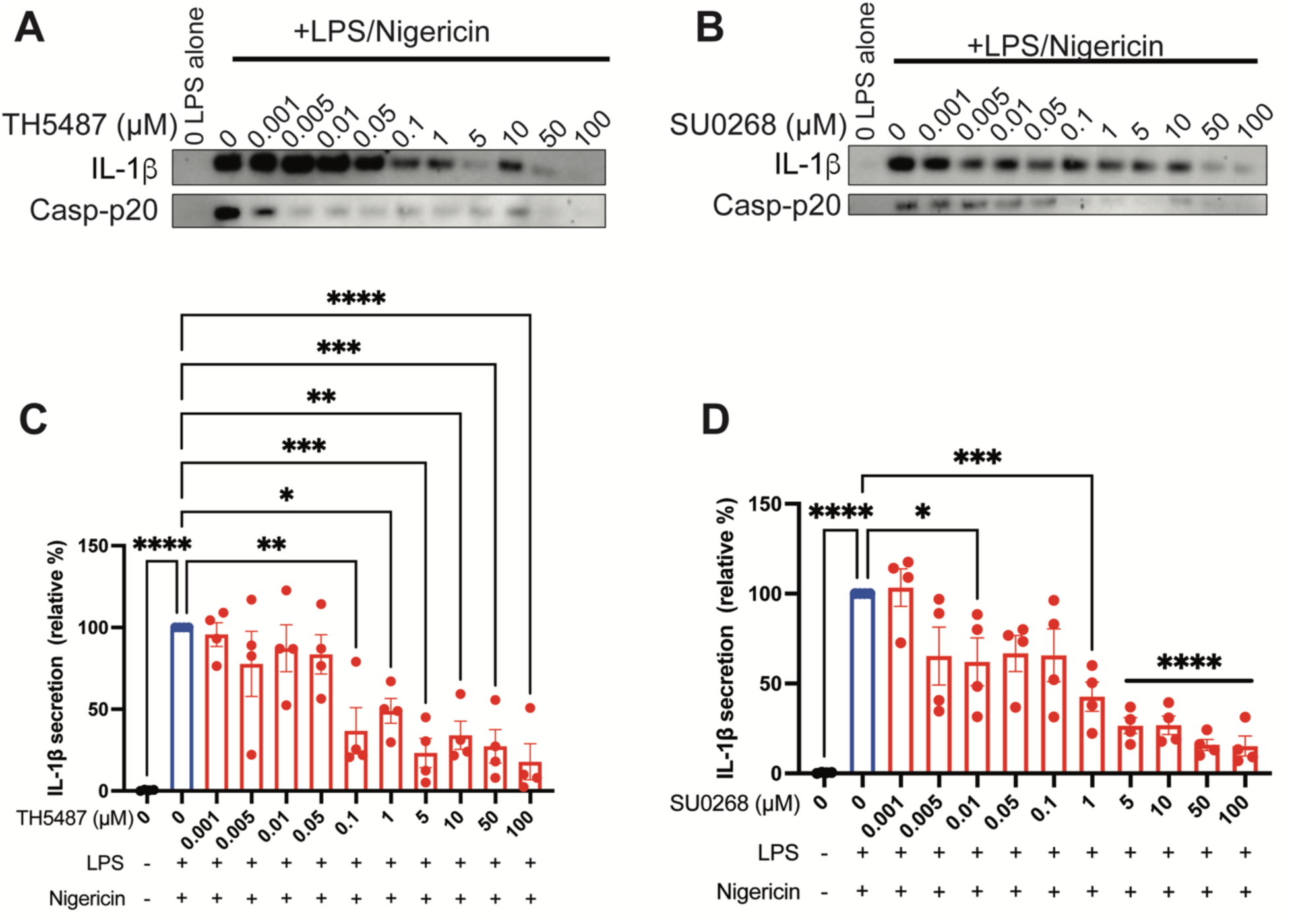
IL-1β section is inhibited by TH5487 and SU0268 in immortalized human THP1s: Immortalized human THP1 cells were activated with LPS and Nigericin and treated with TH5487 or SU0268 at 0.001-100 µM. (A-B) IL-1β release was visualized by western. (C-D) IL-1β release was quantified by relative band intensity. For both graphs, error bars signify the mean +/- SEM. The data was analyzed by one-way ANOVA with N=3-4. p****<0.0001, p***<0.001, p**<0.01, p*<0.05.

**Fig. S2:**
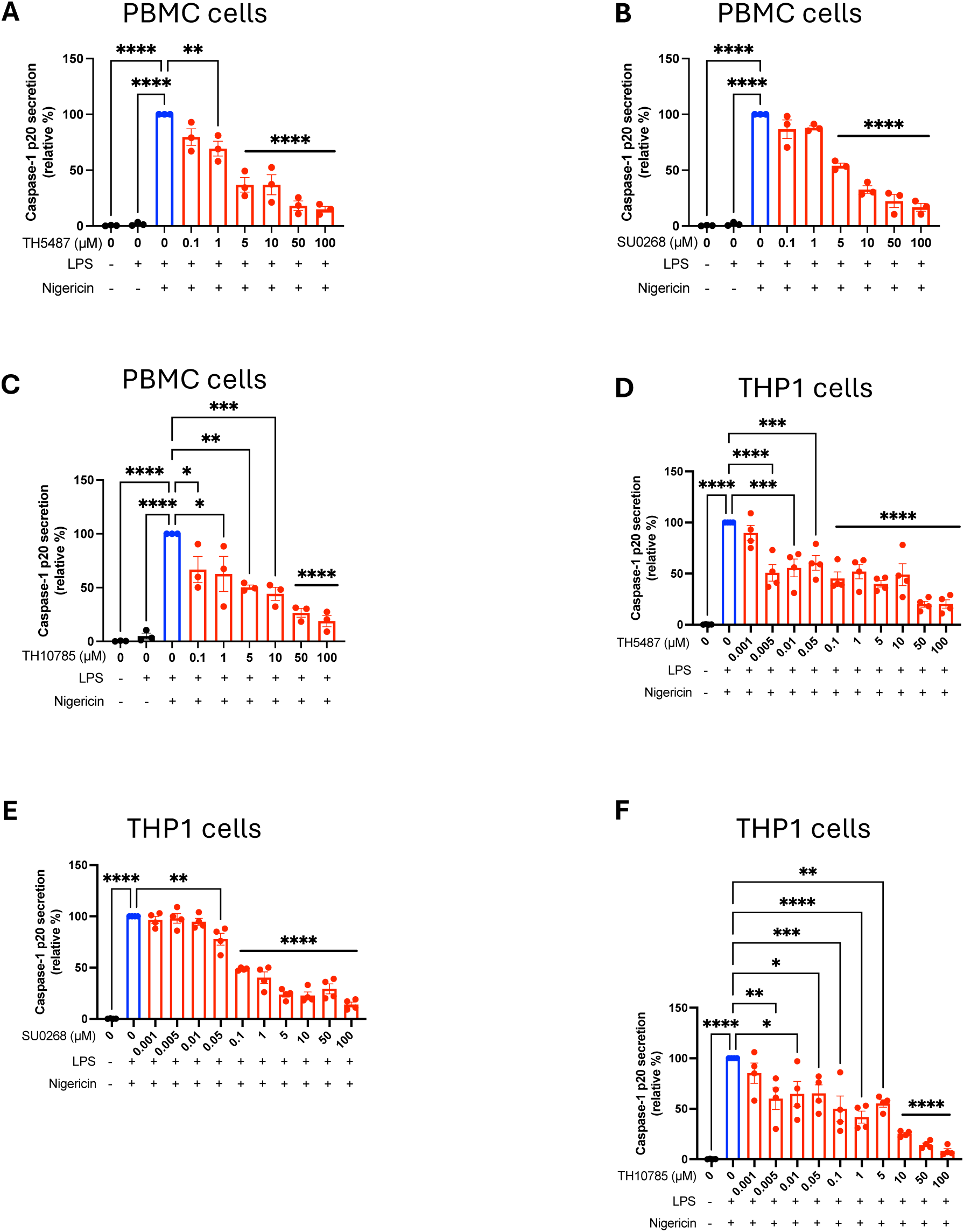
Caspase-1 p-20 section is inhibited by TH5487, SU0268, and TH10785 in primary human PMBCs and immortalized human THP1s. (A-C) Primary human PBMCs were activated with LPS and Nigericin and treated with TH5487, SU0268, or TH10785 at 0.1-100 µM. Casp-p20 release was visualized by western (Fig.1 & 6) and quantified by relative band intensity. (D-F) Immortalized human THP1 cells were activated with LPS and Nigericin and treated with TH5487, SU0268, or TH10785 at 0.001-100 µM. Casp-p20 release was visualized by western (Fig.S1 & S14) and quantified by relative band intensity. For all graphs, error bars signify the mean +/- SEM. The data was analyzed by one-way ANOVA with N=3-4. p****<0.0001, p***<0.001, p**<0.01, p*<0.05.

**Fig. S3:**
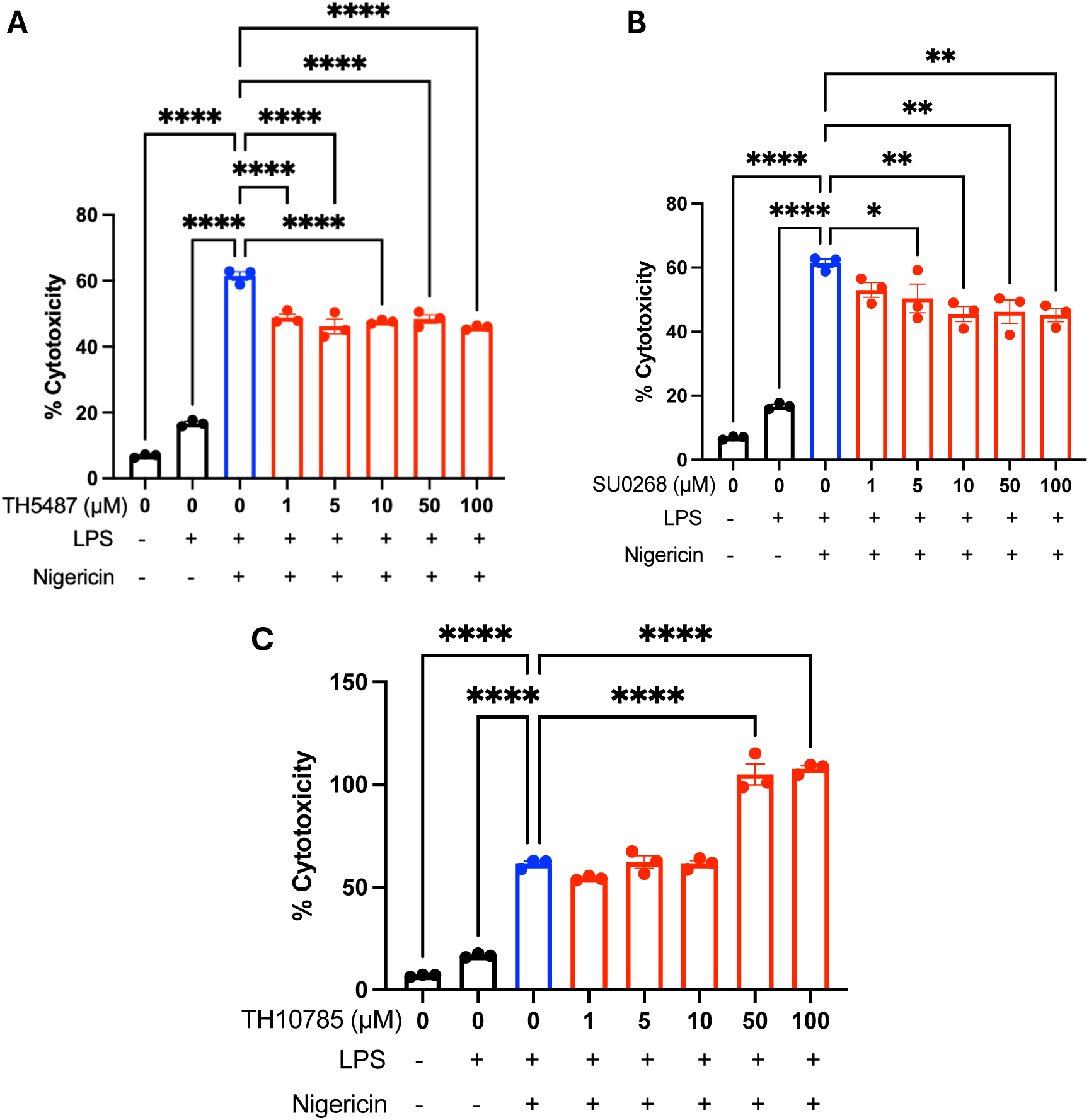
No drugs induce cytotoxicity between 1 and 10 µM. Primary human PBMCs were activated with LPS and Nigericin and treated with small molecules from 1-100 µM. The amount of cytotoxicity was assessed by CyTox 96 fluorescence. A) TH5487 reduced inflammasome-related cytotoxicity from 10-100 µM. B) SU0268 reduced inflammasome-related cytotoxicity from 5-100 µM. C) TH10785 increased cytotoxicity at 50 and 100 µM. For all graphs, error bars signify the mean +/- SEM. The data was analyzed by one-way ANOVA with N=3. p****<0.0001, p**<0.01, p*<0.05.

**Fig. S4:**
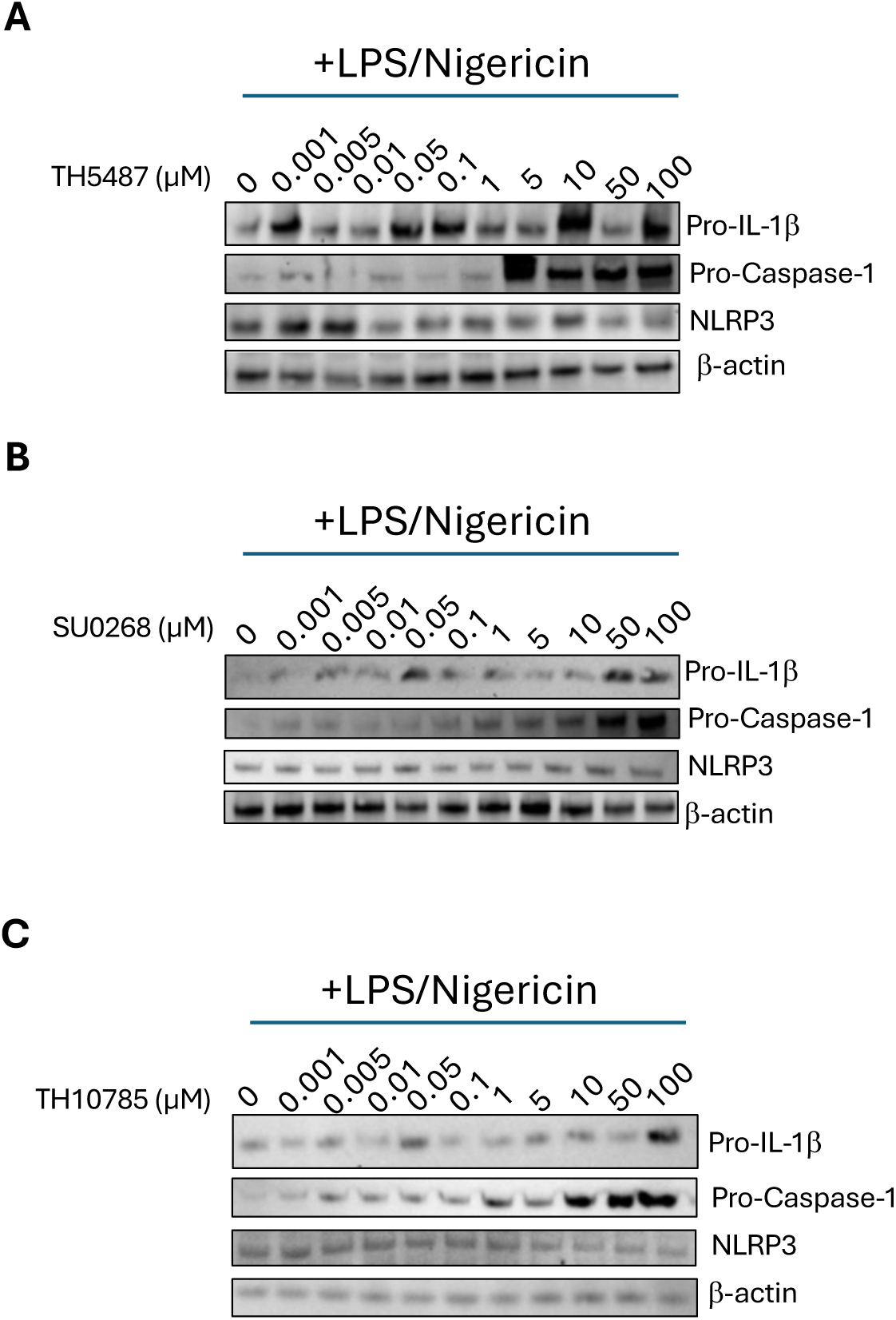
Inhibitors increased the amount of Pro-IL-1β and Pro-Caspase-1 retained in the cell in human THP1 cells, matching the decrease in secretion of mature IL-1β and Casp-p20. Immortalized human THP1 cells were activated with LPS and Nigericin and treated with (A) TH5487, (B) SU0268, or (C) TH10785 at 0.001-100 µM. Pro-IL-1β, Pro-Caspase-1, NLRP3, and β-actin expression were evaluated by western blot.

**Fig. S5.**
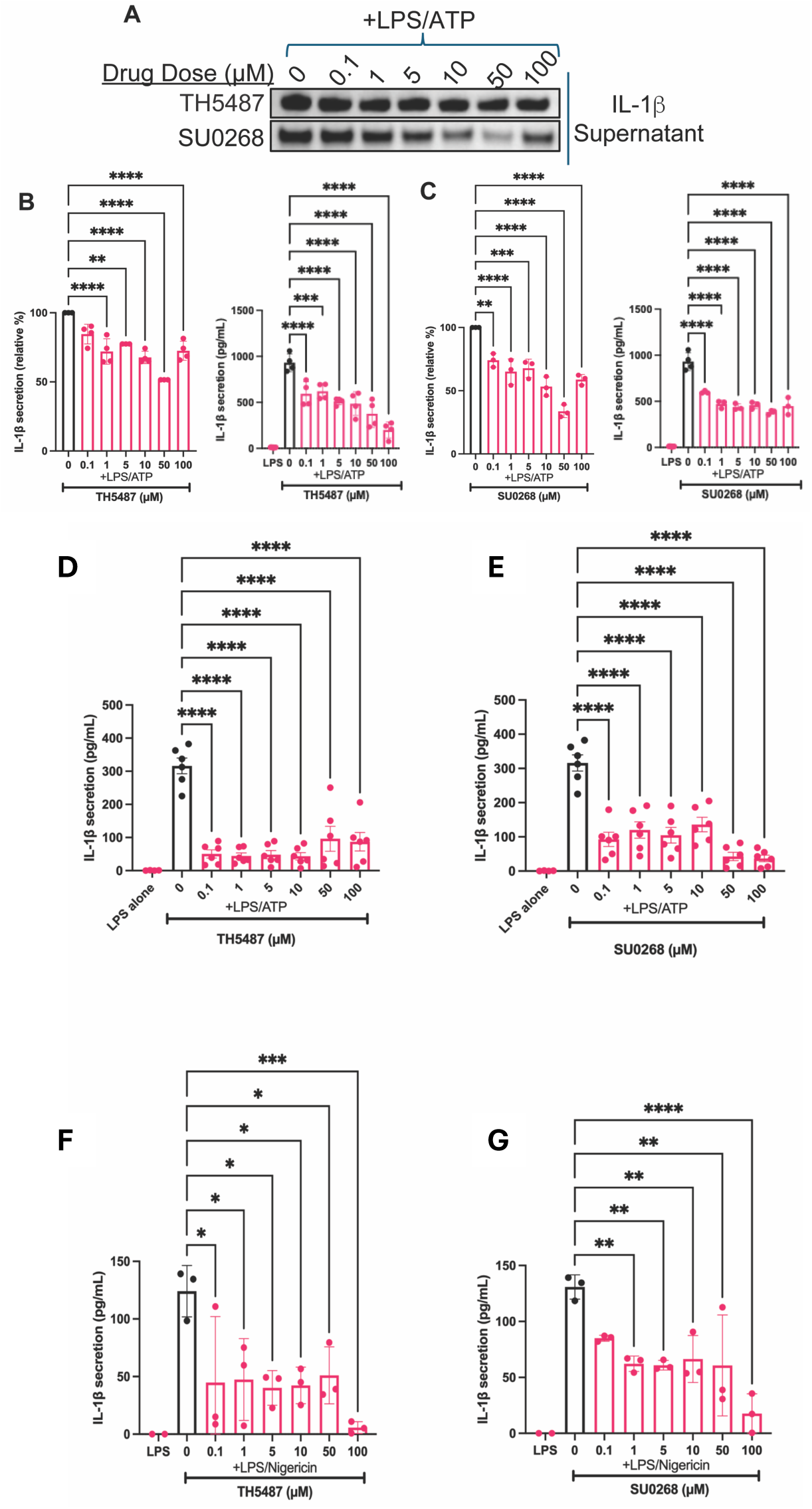
TH5487 and SU0268 inhibit inflammasome activity in human THP1s and mouse iBMDMs activated with either LPS and ATP or LPS and Nigericin. A) THP1 cells activated with LPS and ATP were treated with either TH5487 or SU0268 at 0.1-100 µM. Relative IL-1β release was visualized by western blot. (B-C) Quantification of (A) and confirmation by ELISA for TH5487 data D) Mouse iBMDMs were primed and activated with LPS and ATP and treated with 0.1-100 µM TH5487. The amount of IL-1β secreted into the media was measured using an IL-1β ELISA. E) Mouse iBMDMs were primed and activated with LPS and ATP and treated with 0.1-100 µM SU0268. The amount of IL-1β secreted into the media was measured using an IL-1β ELISA. F) Mouse iBMDMs were primed and activated with LPS and Nigericin and treated with 0.1-100 µM TH5487. The amount of IL-1β secreted into the media was measured using an IL-1β ELISA. G) Mouse iBMDMs were primed and activated with LPS and Nigericin and treated with 0.1-100 µM SU0268. The amount of IL-1β secreted into the media was measured using an IL-1β ELISA. For all graphs, error bars signify the mean +/- SEM. The data was analyzed by one-way ANOVA with N=3-6. p****<0.0001, p**<0.01, p*<0.05.

**Fig. S6.**
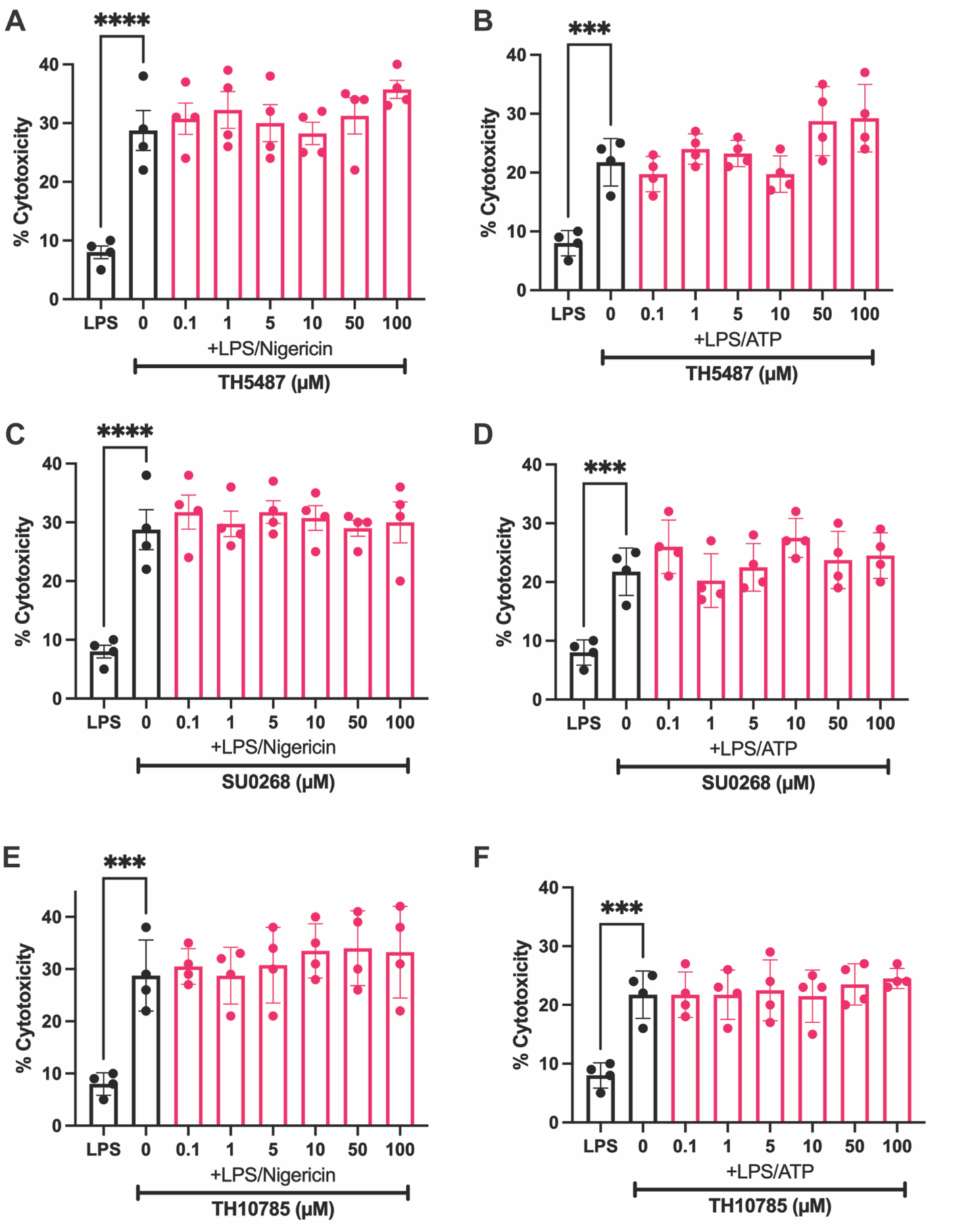
None of the hOGG1 small molecules caused significant toxicity at doses from 0.1-100 µM in human THP1 cells activated with either LPS/ATP or LPS/Nigericin. A) Human monocyte THP1 cells were activated with LPS and Nigericin and treated with TH5487 from 0.1-100 µM. The cell death was quantified by the presence of lactose dehydrogenase (LDH) in the supernatant of treated cells. B) Human monocyte THP1 cells were activated with LPS and ATP and treated with TH5487 from 0.1-100 µM. The cell death was quantified by the presence of lactose dehydrogenase (LDH) in the supernatant of treated cells. C) Human monocyte THP1 cells were activated with LPS and Nigericin and treated with SU0268 from 0.1-100 µM. The cell death was quantified by the presence of lactose dehydrogenase (LDH) in the supernatant of treated cells. D) Human monocyte THP1 cells were activated with LPS and ATP and treated with SU0268 from 0.1-100 µM. The cell death was quantified by the presence of lactose dehydrogenase (LDH) in the supernatant of treated cells. E) Human monocyte THP1 cells were activated with LPS and Nigericin and treated with TH10785 from 0.1-100 µM. The cell death was quantified by the presence of lactose dehydrogenase (LDH) in the supernatant of treated cells. F) Human monocyte THP1 cells were activated with LPS and ATP and treated with TH10785 from 0.1-100 µM. The cell death was quantified by the presence of lactose dehydrogenase (LDH) in the supernatant of treated cells. For all graphs, error bars signify the mean +/- SEM. The data was analyzed by one-way ANOVA with N=3-6. p****<0.0001, p***<0.001

**Fig. S7:**
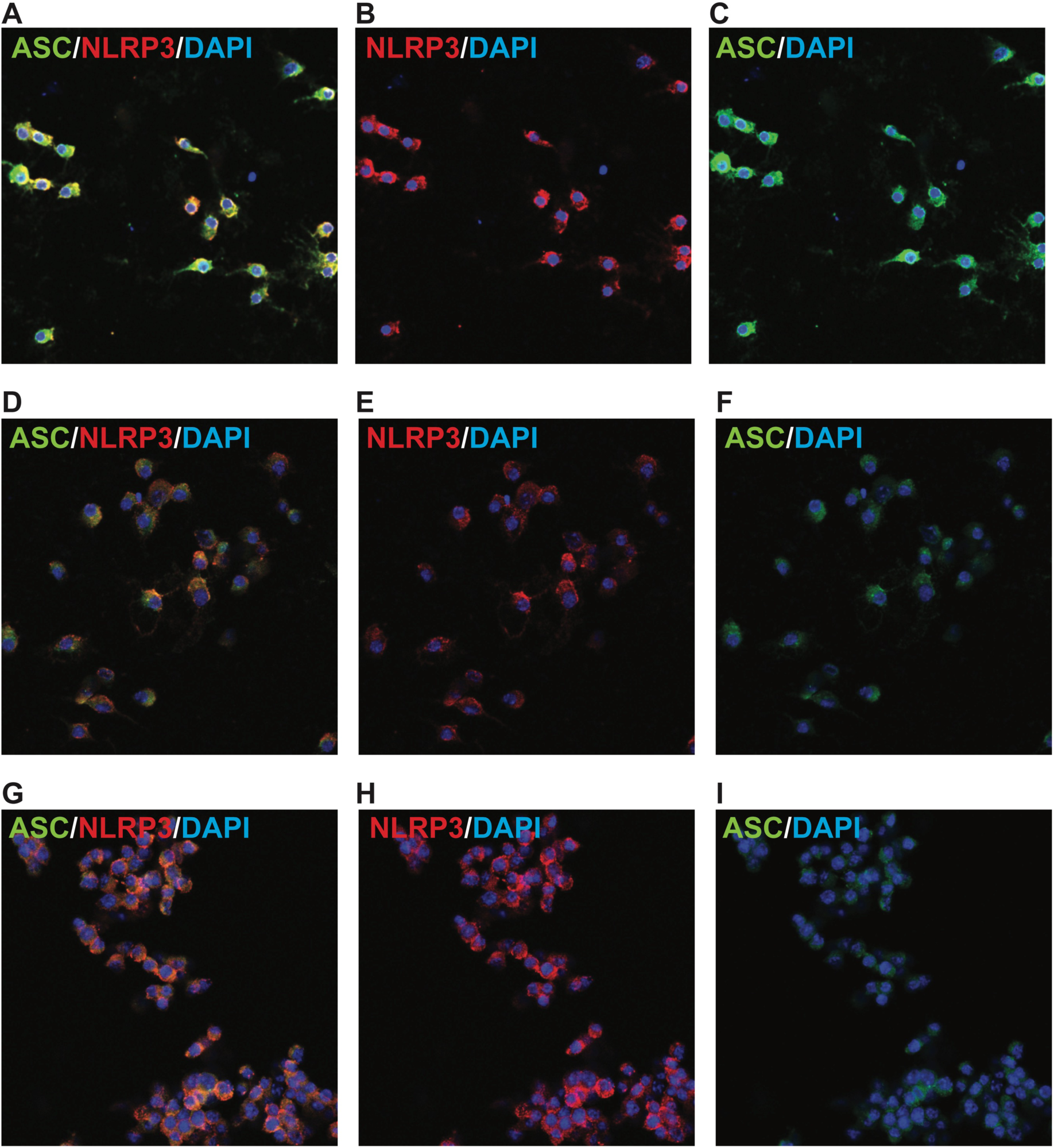
Full scale images from immunofluorescence data in Fig.3. (A-C) iBMDMs were treated with LPS and Nigericin and 0 µM SU0268 then probed for ASC (A), NLRP3 (B), or both (C). (D-F) iBMDMs were treated with LPS and Nigericin and 50 µM SU0268 then probed for ASC (D), NLRP3 (E), or both (F). (G-I) iBMDMs were treated with LPS and Nigericin and 100 µM SU0268 then probed for ASC (G), NLRP3 (H), or both (I).

**Fig. S8:**
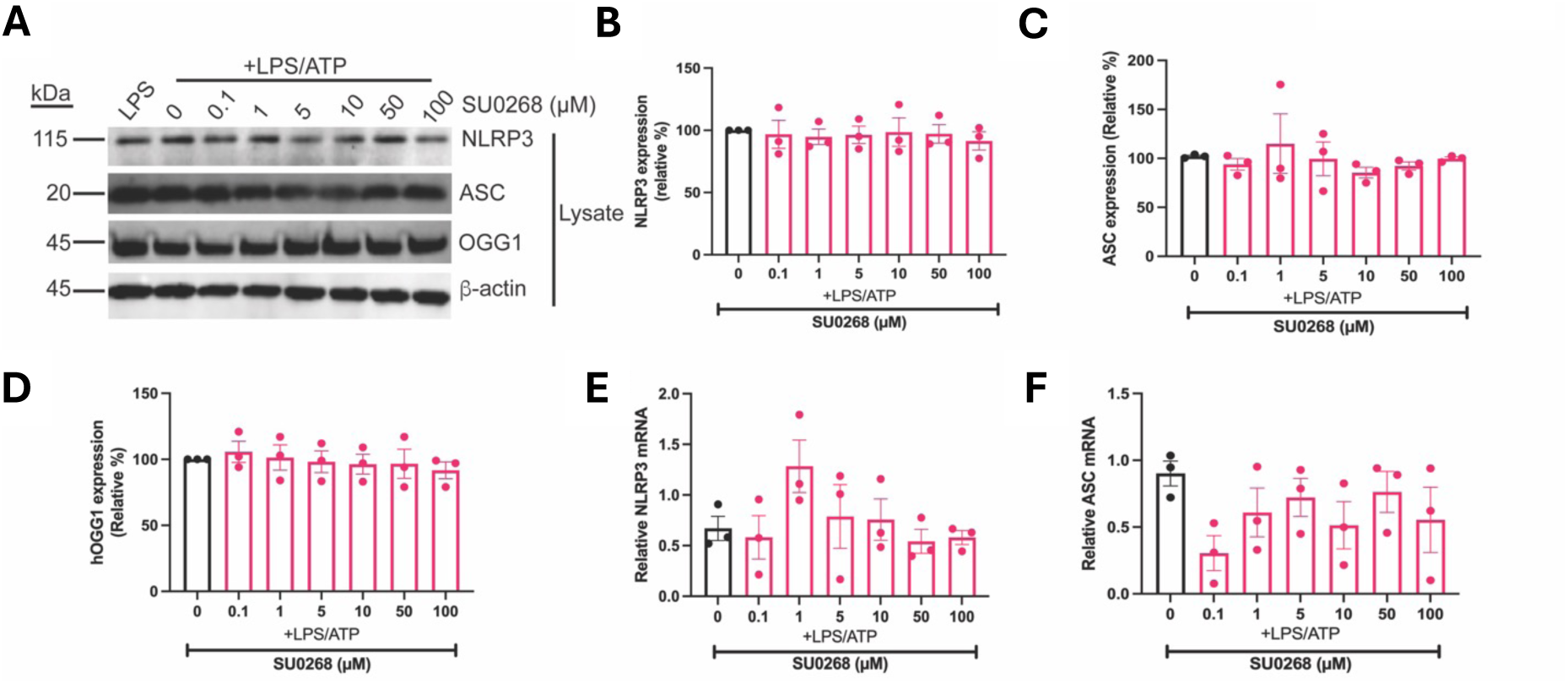
ASC speck and inflammasome assembly inhibition do not effect protein expression. A) BMDMS were primed with LPS and activated with ATP, and treated with SU0268. The lysate of the cells were probed with antibodies for NLRP3, ASC, OGG1, and b-actin and visualized by western blot. The band intensities of western blots against NLRP3 (B), ASC (C), and OGG1 (D) were quantified and compared to the untreated activated control. RNA was isolated from treated cells and RT-PCR followed by qPCR was performed to qwuantify mRNA expression for either NLRP3 (E) or ASC (F). For all graphs, error bars signify the mean +/- SEM. The data was analyzed by one-way ANOVA with N=3.

**Fig. S9:**
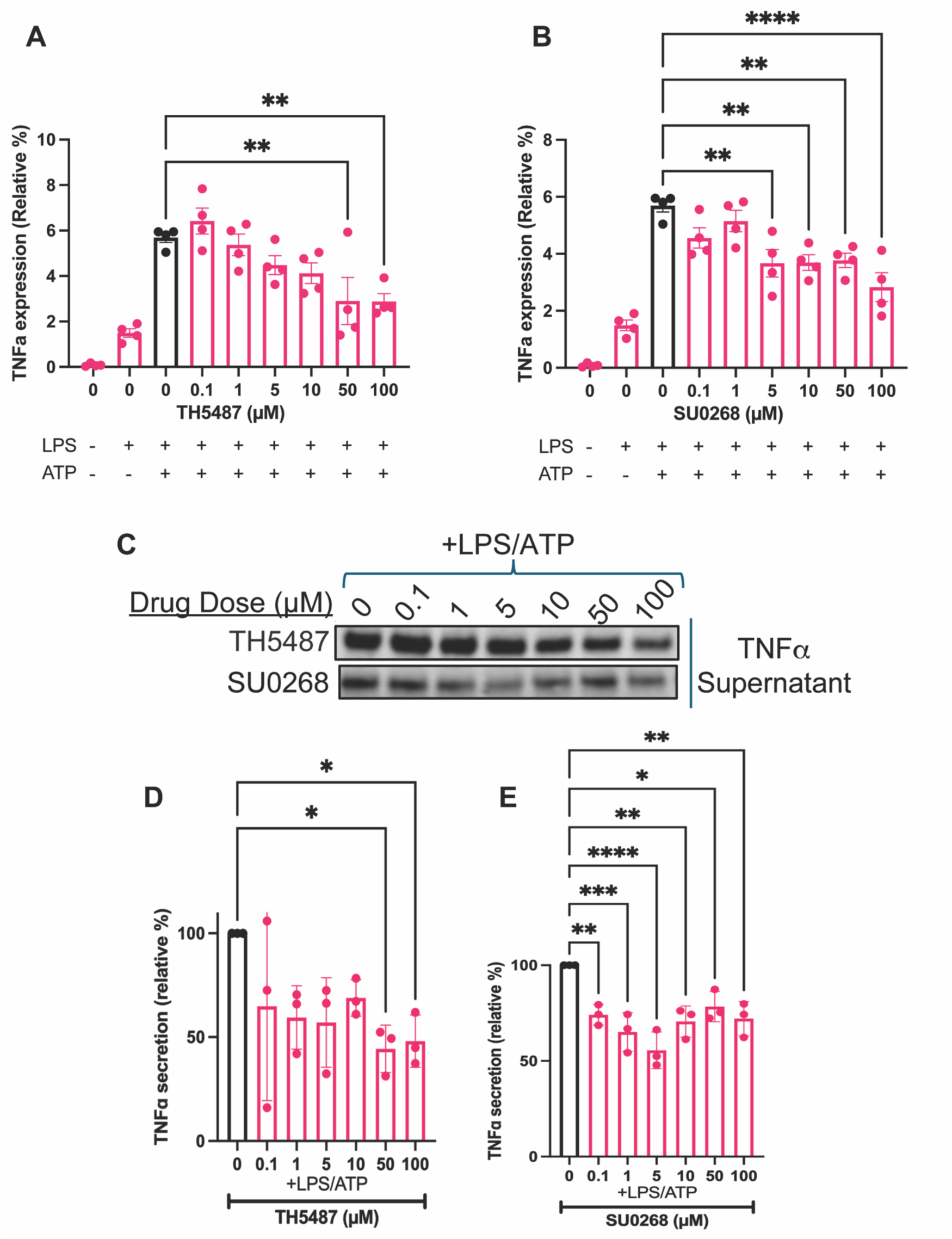
TH5487 and SU0268 inhibit inflammatory signaling activity in human THP1 cells. THP1 cells activated with LPS and ATP were treated with either TH5487 or SU0268 at 0.1-100 µM. A) The expression of TNF-α in cells treated with TH5487 was evaluated by qPCR. B) The expression of TNF-α in cells treated with SU0268 was evaluated by qPCR C) Relative TNF-α release was visualized by western blot. D) quantification of (C) for TH5487 data. E) Quantification of (C) for SU0268 data. For all graphs, error bars signify the mean +/- SEM. The data was analyzed by one-way ANOVA with N=3-4. p****<0.0001, p***<0.001, p**<0.01, p*<0.05.

**Fig. S10:**
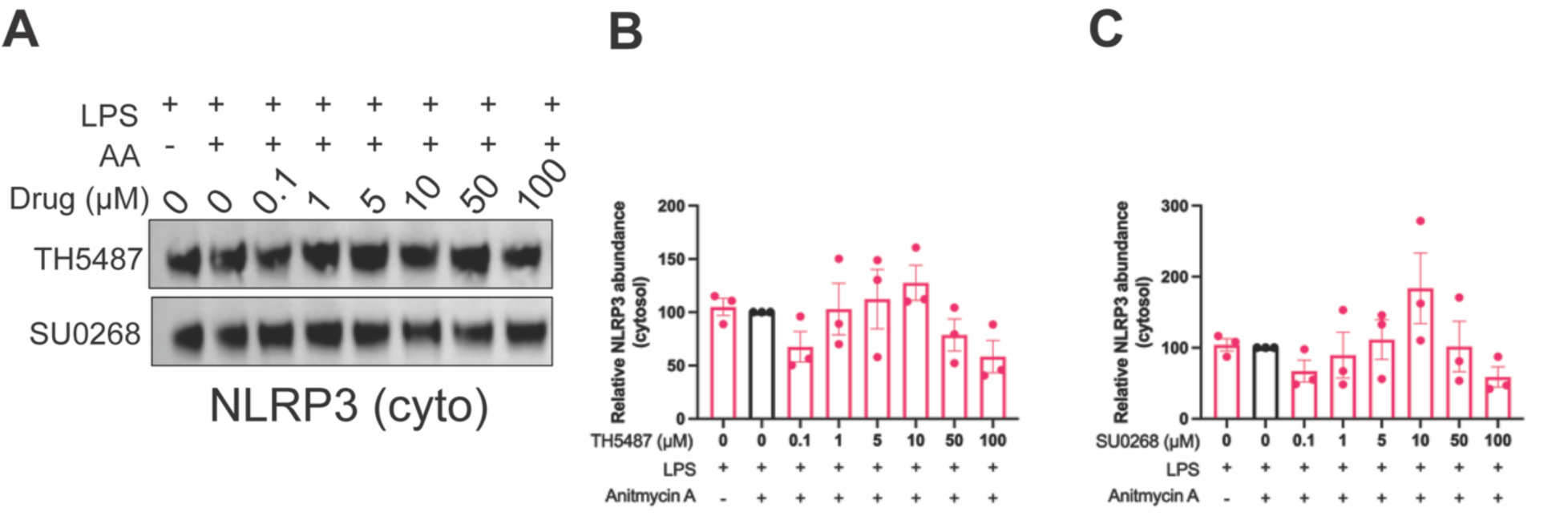
During mitochondrial dysfunction and subsequent treatment with small molecules, the majority of NLRP3 remains in the cytosol. A) THP1 cells activated with LPS and Antimycin A were treated with either TH5487 or SU0268 at 0.1-100 µM. The cytosolic fraction was isolated and the relative amount of NLRP3 in the cytosolic fraction was visualized by western blot. B) quantification of (A) for TH5487. C) Quantification of (A) for SU0268. For all graphs, error bars signify the mean +/- SEM. The data was analyzed by one-way ANOVA with N=3-4.

**Fig. S11:**
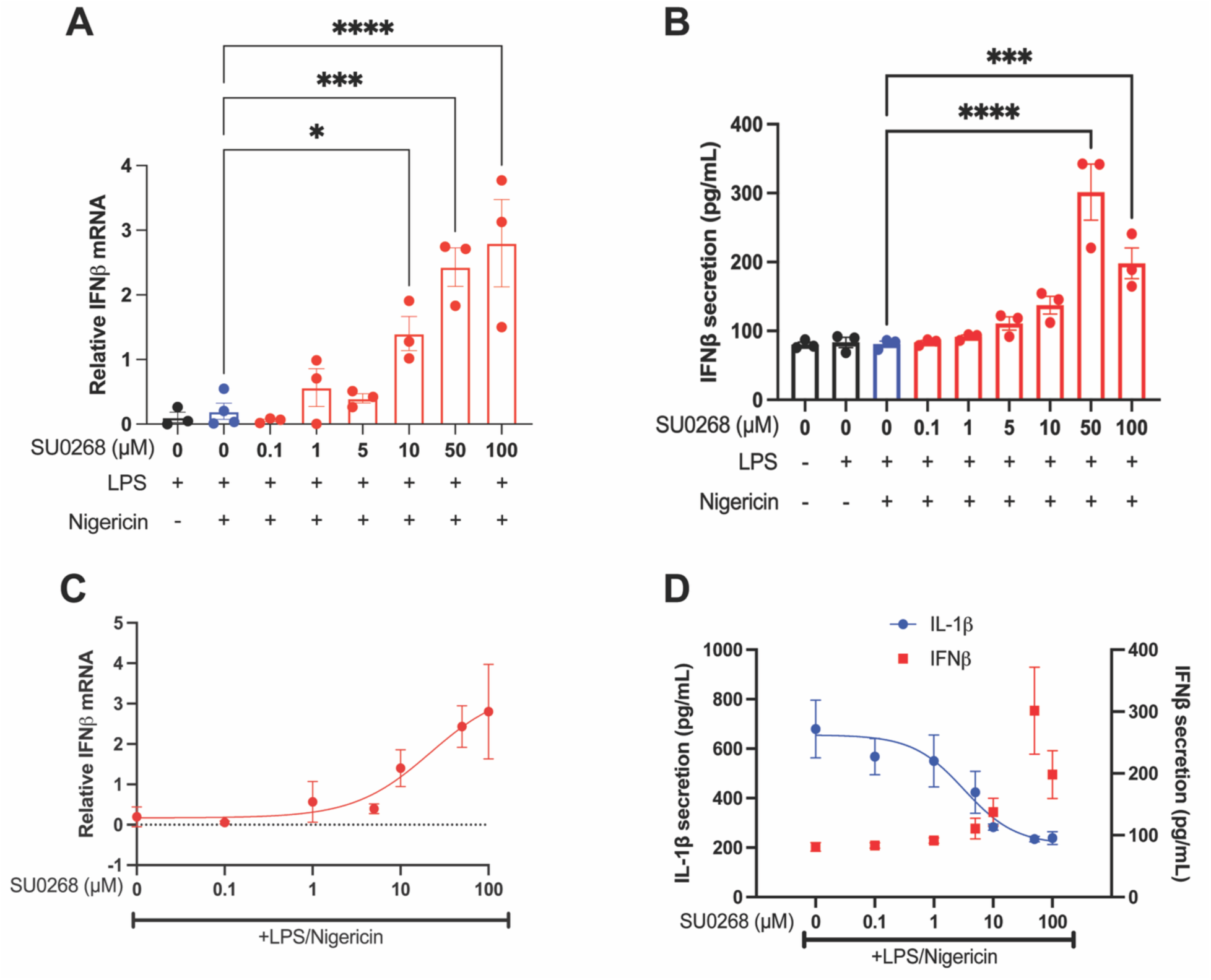
SU0268 inhibits NLRP3 activity and activates the cGAS-STING pathway in primary human PBCMs. Primary human PBMCs from healthy donors were activated with LPS and Nigericin and treated with SU0268 at 0.1-100 µM. A) IFN-β expression was quantified by qPCR. B) IFN-β secretion was quantified by ELISA. C) The EC50 of TH5487 for IFN-β was found using non-linear regression at 21.92 µM with R^2^=0.81. D) IL-1β and IFN-β secretion was quantified by ELISA and compared using non-linear regression showing inversely related trends of expression. IC50_SU0268_: 3.25 µM, R^2^_SU0268_: 0.8209. For bar graphs, error bars signify the mean +/- SEM. The data was analyzed by one-way ANOVA with N=3-4. p****<0.0001, p***<0.001, p*<0.05. For the line graph, error bars signify mean +/- SD and the data was analyzed using non-linear regression.

**Fig. S12:**
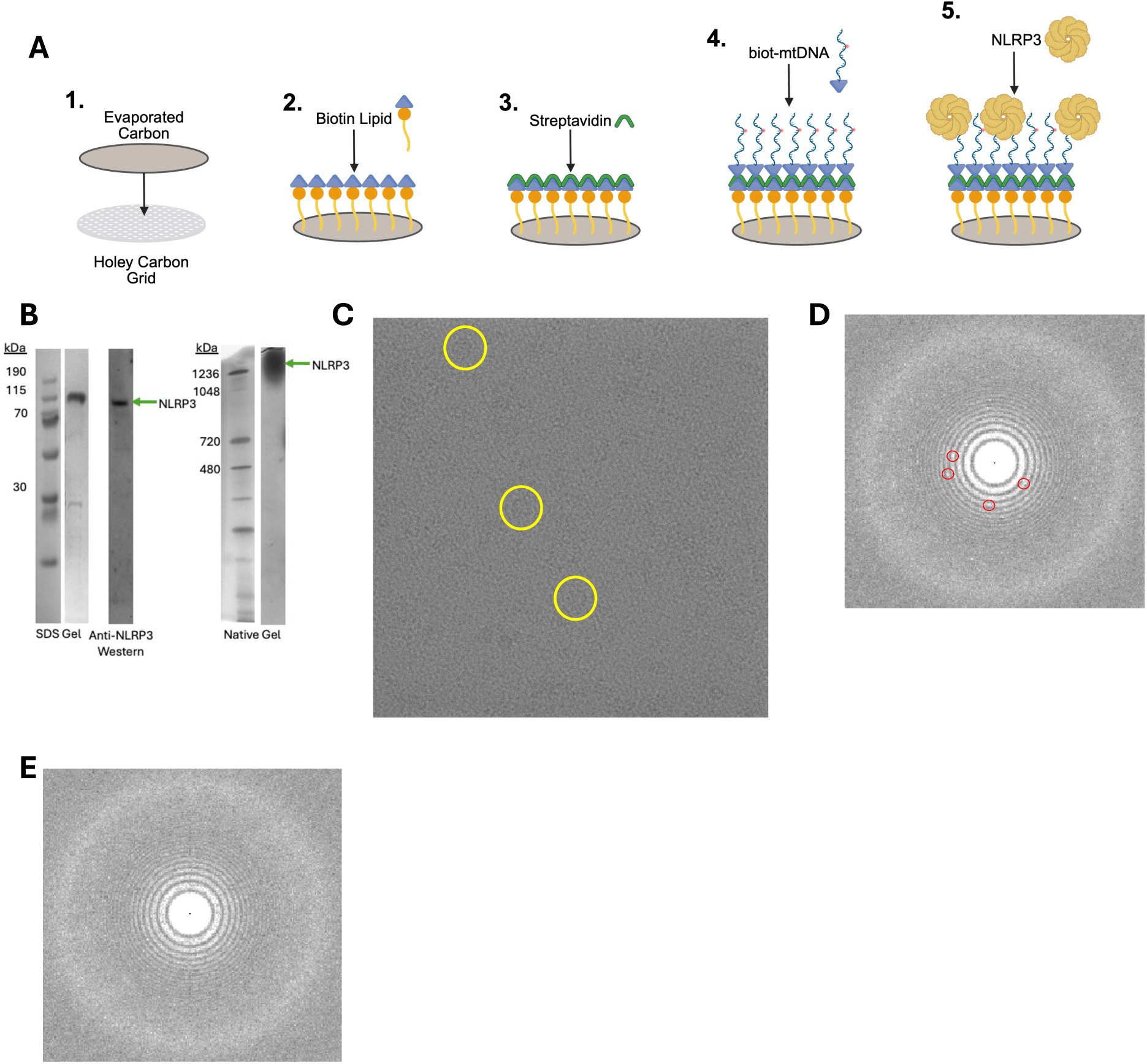
NLRP3 purified in the presence of Th5487 can be visualized on streptavidin-affinity grids using cryoEM. A) Schematic of streptavidin grid preparation workflow B) Full length human NLRP3 was expressed in Expi293 cells and purified by affinity and size exclusion chromatography. On an SDS Page gel and western blot, a band associated with NLRP3 is shown at the expected size of ∼120 kDa. On a native gel, a band associated with an NLRP3 decamer is shown at the expected size of ∼1200 kDa. C) Example cryoEM micrograph before streptavidin lattice removal. Yellow circles show NLRP3 particles bound to the streptavidin/biot-DNA layer. D) FFT of un-subtracted micrograph. Red circles show speckle contamination caused by streptavidin lattice. E) FFT of lattice-subtracted micrograph with no speckle contamination.

**Fig. S13.**
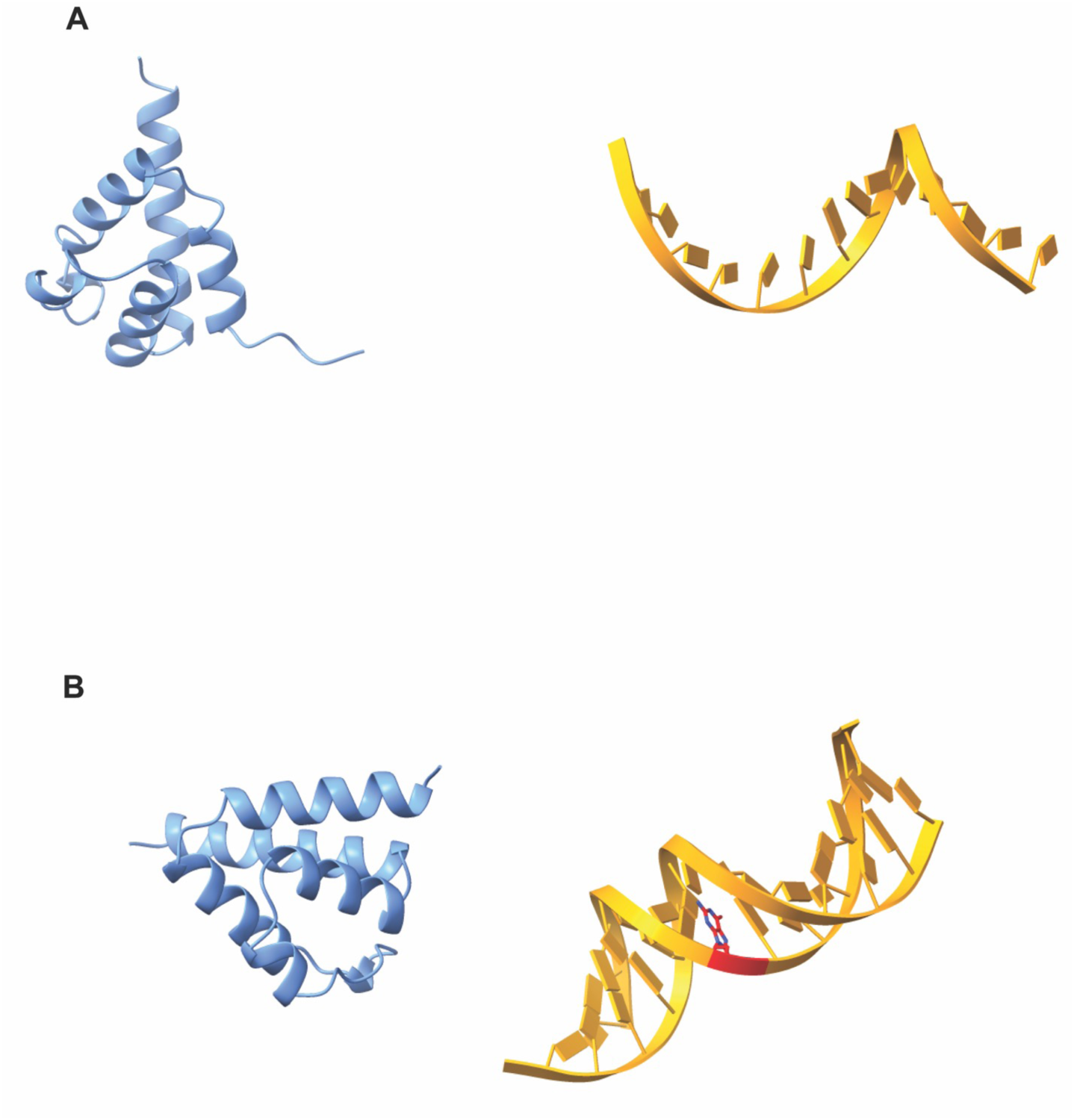
AlphaFold does not predict the NLRP3 pyrin domain will interact with single stranded non-oxidized DNA or double stranded oxidized DNA. A) AlphaFold model of the NLRP3 pyrin domain (residues 1-85, blue) in the presence of single stranded non oxidized DNA (orange). B) AlphaFold model of the NLRP3 pyrin domain (residues 1-85, blue) in the presence of double stranded (orange) oxidized DNA (red, from PDBID: 1EBM).

**Fig. S14:**
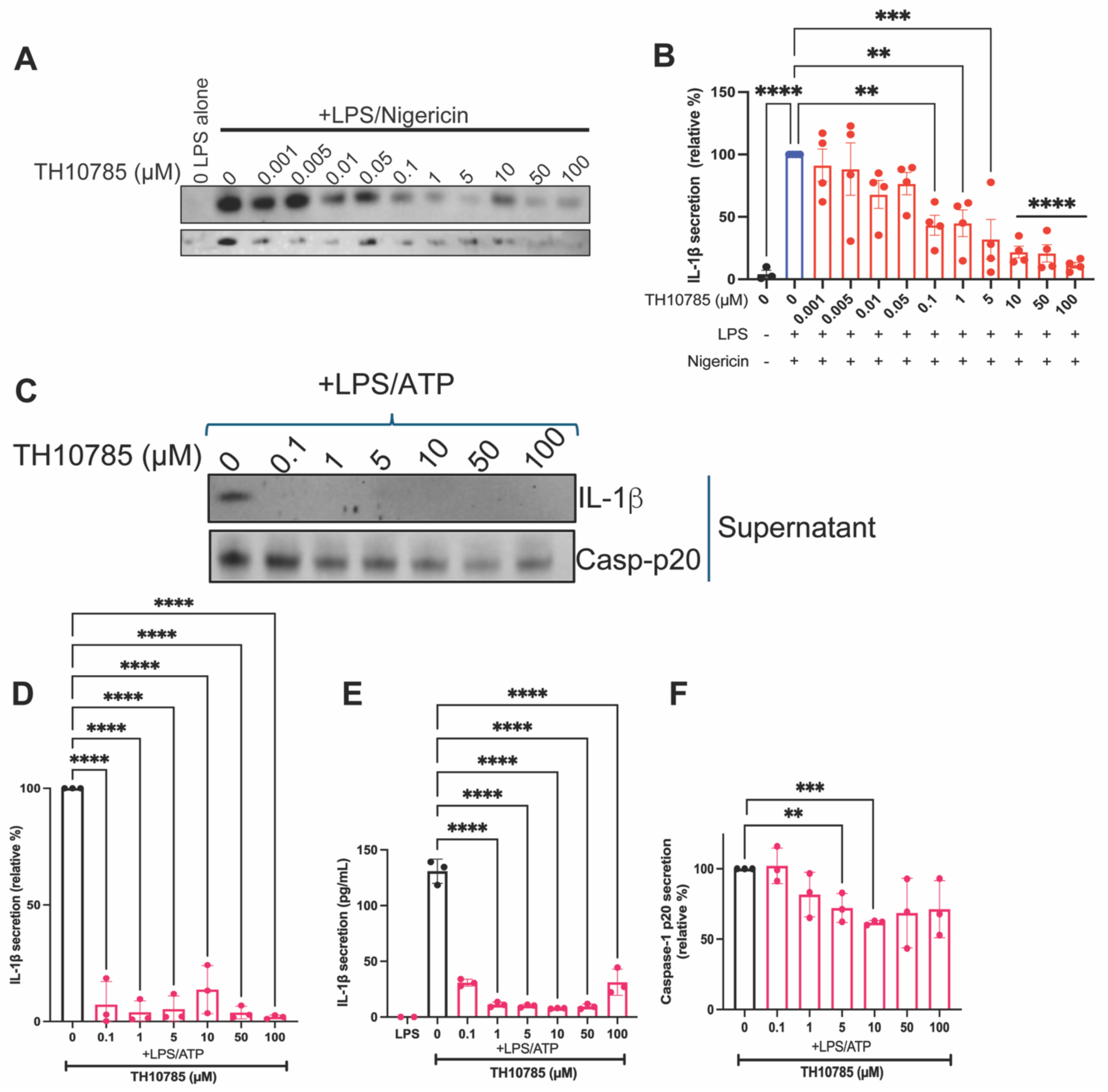
TH10785 inhibits inflammasome activation in mouse macrophages. Immortalized human THP1 cells were activated with LPS and Nigericin and treated with TH10785 at 0.001-100 µM. (A) IL-1β release was visualized by western. (B) IL-1β release was quantified by relative band intensity.C) Mouse iBMDMs activated with LPS and ATP were treated with TH10785 at 0.1-100 µM. The amounts of secreted IL-1β and Casp-p20 were visualized by western blot. (D/E) The amount of secreted IL-1β was quantified both by western blot (D) and ELISA (E). (F) The amount of secreted Casp-p20 was quantified by western blot. For all graphs, error bars signify the mean +/- SEM. The data was analyzed by one-way ANOVA with N=3. p****<0.0001, p***<0.001, p**<0.01.

**Table S1:**
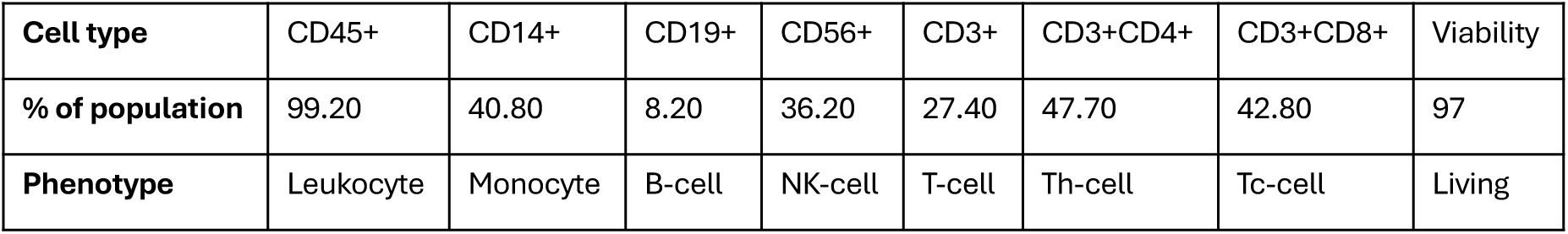
Morphology of primary human PBMC cell types.

|  |  |  |  |  |  |  |  |  |
| --- | --- | --- | --- | --- | --- | --- | --- | --- |
| **Cell type** | CD45+ | CD14+ | CD19+ | CD56+ | CD3+ | CD3+CD4+ | CD3+CD8+ | Viability |
| **% of population** | 99.20 | 40.80 | 8.20 | 36.20 | 27.40 | 47.70 | 42.80 | 97 |
| **Phenotype** | Leukocyte | Monocyte | B-cell | NK-cell | T-cell | Th-cell | Tc-cell | Living |

**Table S2.** Measured distanced between the NLRP3 pyrin structure or the NLRP3 pyrin domain model and hOGG1 bound to TH10785 presented in Fig.5. The locations of active site amino acids in hOGG1 bound to TH10785 (PDB: 7AYY, 249-325) were compared to that of similar amino acids in the NLRP3 pyrin domain (PDB: 7PZC, 1-91) (third from right column) or the SWISS-MODEL generated model of the NLRP3 pyrin domain (far left column).

| WT NLRP3: AA codes | hOGG1: AA codes | Distance (Å)<br>hOGG1<br>TH10785/WT | Distance (Å)<br>hOGG1<br>TH10785/Model |
| --- | --- | --- | --- |
| Lys 2 | Glu/Lys 249 |  | 0.632 |
| Ala 4 | Ala 251 | 10.556 | 0.439 |
| Cys 8 | Cys 255 | 7.906 | 0.373 |
| Ala 11 | Ala 258 | 16.110 | 0.286 |
| Asp 16 | Asp 268 | 3.571 | 17.863 |
| Asp 19 | Asp 268 | 7.782 | 13.315 |
| Val 20 | Val 267 | 13.134 | 6.447 |
| Asp 21 | Asp 268 | 8.048 | 7.031 |
| His 28 | His 270 | 10.53 | 5.463 |
| Asp 31 | Asp 278 | 3.378 | 4.847 |
| Tyr 32 | Tyr 279 | 6.641 | 4.436 |
| Pro 34 | Pro 283 | 24.239 | 2.473 |
| Pro 42 | Pro 291 | 17.71 | 11.260 |
| Gln 45 | Gln 294 | 13.188 | 0.634 |
| Thr 46 | Thr 295 | 9.240 | 0.637 |
| Leu 54 | Leu 299 | 0.420 | 0.378 |
| Gly 63 | Gly 308 | 0.528 | 0.101 |
| Trp 68 | Trp 313 | 3.621 | 0.098 |
| Ala 69 | Ala 314 | 1.392 | 0.176 |
| Ala 71 | Ala 316 | 4.595 | 0.571 |
| Val 72 | Val 317 | 3.926 | 0.793 |
| Phe 75 | Phe 319 | 1.991 | 0.649 |
| Ala 77 | Ala 321 | 3.995 | 0.949 |
| Arg 80 | Arg 324 | 9.094 | 1.279 |

## METHODS

### Primary human PBMC inflammasome activation in the presence of TH5487, SU0268, and TH10785 Immortalized Human THP1 Cell Culture

Immortalized wild-type and NLRP3 knockout human THP1 cells were purchased from InvivoGen (THP1-Null and THP1-KO-NLRP3, respectively). Frozen vials were thawed in a water bath and then resuspended in 50 mL of culture media (RPMI no phenol red (Gibco), 10% heat-inactivated FBS (Sigma), 1x Penicillin-Streptomycin-Glutamine (Gibco), 1x non-essential amino acids (Gibco), 1x Sodium Pyruvate (Gibco)). The cells were spun down at 300 x g for 5 minutes at 4°C, the pellet was resuspended in 5 mL of culture media. The cells were counted and resuspended as described above, doubling every 24-48 hours. The viability and concentration was checked by measuring the concentration on a 1 mL aliquot of the cells using Trypan Blue as described above, and expanded accordingly.

### THP1 NLRP3 Inflammasome Activation in the presence of TH5487, SU0268, and TH10785

The viability and concentration of a culture of immortalized mouse macrophages were checked as described above. On day 0, cells at viability >95% and 0.5*106 cells/mL were split into 6-well TC-treated plates (Corning) with 2 mL of cells per well and allowed to double overnight. On day 1, 2 µL/mL of 500x lipopolysaccharide (Thermo Fisher) was added to each well for 16 hours for a final concentration of 400 ng/mL. On day 2, LPS-only wells were harvested, and drugs TH5487 (Selleck Chemicals), SU0268 (MedChemExpress), and/or TH10785 were serial diluted in DMSO (Fisher Scientific) such that the addition of any concentration of inhibitor was 1% of the final volume of cells. Then the inhibitors were added at concentrations ranging from 0.1-100 µM for 1 hour. Next, 20 µM Nigericin (Sigma) or 4 mM ATP was added to each well for 1 hour. Cells and supernatant fractions could then be isolated for viability and western blot analysis. The supernatant and cells were separated by spinning at 300 x g for 5 minutes at 4°C. The supernatant fraction was removed from the cell pellet and clarified by spinning at 3000 xg for 5 minutes using 0.22 µm spin filters (Corning). The cell pellet was resuspended in 1 mL ice-cold PBS and re-pelleted by spinning at 1000 x g for 5 minutes at 4°C. The PBS was removed from the pellet and the cells were lysed with 300 µL of RIPA buffer (Boston BioProducts) supplemented with an EDTA-free protease/phosphatase inhibitor cocktail (Roche) by rotating for 5 minutes at 4°C. The lysed cells were spun at 14,000 x g for 15 minutes. 200 µL of the clarified lysate was removed and saved for analysis. The protein concentration of each fraction (supernatant and whole cell) was evaluated using a Bradford Assay (BioRad) and western blots were run to quantify protein expression

### Immortalized Bone Marrow-Derived Mouse Macrophage Cell Culture

Immortalized wild-type bone marrow derived mouse macrophages were generously provided to us by Michael Karin at the University of California, San Diego. A frozen vial of cells at 1*107 was thawed in a water bath and then resuspended in 50 mL of culture media (DMEM (Thermo Fisher) supplement with 10% heat-inactivated FBS (Sigma) and 1% Penicillin-Streptomycin (Thermo Fisher)). The cells were spun down at 300 x g for 5 minutes at 4°C, the pellet was resuspended in 5 mL of culture media. The cells were counted by diluting 10 µL of cells with 10 µL of Trypan Blue Stain (Thermo Fisher), loading that dilution onto a CountessTM Cell Counting Chamber Slide (Thermo Fisher), and evaluated using a CountessTM 3 FL Automated Cell Counter (Thermo Fisher). Cells were then further diluted in culture media such that the final concentration was 0.2*106/mL and plated onto 175cm2 culture flasks (Corning) where they were grown at 37°C and 5% humidity. Cells typically doubled in 24-48 hours, where they were then scraped from the bottom of the plate using Bio-One Cell Scrapers (Fisher Scientific) and then spun down, counted, and expanded as previously described.

### Macrophage NLRP3 Inflammasome Activation in the presence of TH5487, SU0268, and TH10785

The viability and concentration of a culture of immortalized mouse macrophages were checked as described above. Cells at viability >95% and 0.5*106 cells/mL were split into 6-well TC-treated plates (Corning) with 2 mL of cells per well and allowed to adhere and double overnight. The next day, 2 µL/mL of 500x lipopolysaccharide (Thermo Fisher) was added to each well for 1 hour for a final concentration of 400 ng/µL. Next, LPS-only wells were harvested, and drugs TH5487 (Selleck Chemicals), SU0268 (MedChemExpress), and/or TH10785 were serial diluted in DMSO (Fisher Scientific) such that the addition of any concentration of inhibitor was 1% of the final volume of cells. Then the inhibitors were added at concentrations ranging from 0.1-100 µM for 1 hour. Next, 20 µM Nigericin (Sigma) or 4 mM ATP was added to each well for 1 hour. Cells and supernatant fractions could then be isolated for viability and western blot analysis. To collect samples for western blot analysis, the supernatant was removed from each well and clarified by spinning at 3000 xg for 5 minutes using 0.22 µm spin filters (Corning). The cells left on the plate were washed with ice-cold PBS and then lysed with 300 µL of RIPA buffer (Boston BioProducts) supplemented with an EDTA-free protease/phosphatase inhibitor cocktail (Roche). The lysis took place for 5 minutes rocking at 4°C. The lysed cells were then collected into 1.5 mL tubes and spun at 14,000 x g for 15 minutes. 200 µL of the clarified lysate was removed and saved for western blot analysis. The protein concentration of each fraction (supernatant and whole cell) was evaluated using a Bradford Assay (BioRad) and western blots were run to quantify protein expression.

### THP1 NLRP3 Inflammasome Activation by Antimycin A in the presence of TH5487 and SU0268

The viability and concentration of a culture of immortalized mouse macrophages were checked as described above. On day 0, cells at viability >95% and 0.5*106 cells/mL were split into 6-well TC-treated plates (Corning) with 2 mL of cells per well and allowed to double overnight. On day 1, 2 µL/mL of 500x lipopolysaccharide (Thermo Fisher) was added to each well for 16 hours for a final concentration of 400 ng/mL. On day 2, LPS-only wells were harvested, and drugs TH5487 (Selleck Chemicals) and/or SU0268 (MedChemExpress) were serial diluted in DMSO (Fisher Scientific) such that the addition of any concentration of inhibitor was 1% of the final volume of cells. Then the inhibitors were added at concentrations ranging from 0.1-100 µM for 1 hour. Next, LPS alone wells were harvested, and Antimycin A (ThermoFisher) was resuspended in DMSO and added to each well for a final concentration of 10 µM for two hours. The supernatant and cells were separated by spinning at 300 x g for 5 minutes at 4°C. The supernatant fraction was removed from the cell pellet and clarified by spinning at 3000 xg for 5 minutes using 0.22 µm spin filters (Corning). The cell pellet was resuspended in 1 mL ice-cold PBS and re-pelleted by spinning at 1000 x g for 5 minutes at 4°C. The PBS was removed, and the pellet was further processed to separate the cytosolic, mitochondrial, and whole cell fractions. One-third of the pellet was used for whole cell lysis as described above. The rest of the pellet was resuspended in mitochondrial extraction buffer (220mM mannitol, 70mM sucrose, 20mM HEPES KOH pH 7.5, 1mM EDTA, 2mg/mL BSA) and passed through a 25-G syringe 20 times on ice. Samples were centrifuged at 1000xg for 15min at 4c, and the supernatant was saved in a new tube. These tubes were further centrifuged at 10,000xg for 10min at 4c to collect the mitochondria. The supernatant was saved as the cytosolic fraction, and the pelleted mitochondrial fraction was lysed using the whole cell lysis method. The protein concentration of each fraction (supernatant, whole cell, cytosolic, and mitochondrial) was evaluated using a Bradford Assay (BioRad) and western blots were run to quantify protein expression.

### Protein Quantification by Western Blot

Protein concentration determination and normalized using a Bradford Assay (abcam #ab102535). To evaluate proteins secreted from the cells, the supernatant fractions (Caspase-1 and IL-1β) were run on western blots. To evaluate proteins expressed inside the cell (NLRP3, Caspase-1, FEN1, hOGG1, and Actin), the lysate fraction was run. Samples were diluted with LDS sample loading buffer and reducing agent (Invitrogen), each at a final concentration of 1x. The samples were boiled at 90°C and run on NuPAGE™ 4 to 12%, Bis-Tris 1 mM 15-well mini-gels at 200 V for 30 minutes. Samples were transferred to PVDF membranes, blocked with 2.5% BSA in TBST, and probed with a primary antibody against the specific protein diluted to the manufacturer’s recommendation in 2.5% BSA in TBST. Blots were incubated with an HRP-linked secondary antibody (either mouse, rabbit, or goat depending on the species of the primary) and imaged using the iBright 1500 Imaging system. The intensities of the bands were quantified using the iBright Analysis Software. The intensity values were plotted and analyzed using GraphPad Prism and a one-way ANOVA.

### IL-1β Secretion Quantification by ELISA

Inflammasome activation was measured by an enzyme-linked immunosorbent assay (ELISA) against human or mouse IL-1β, depending on if the samples were from primary human PBMCs, THP1 cells, or iBMDMs, respectively. Both human and mouse IL-1β kits were purchased from Abcam, and used according to the manufacturer’s instructions. Each assay was performed the same, with specific reagents for each kit. Breifly, Standards were reconstituted in Standard Diluent Buffer and serial diluted 5 times from 500-15.6 pg/mL (human) or 6 times from 100-1.56 pg/mL (mouse). A buffer alone blank control was also included for both. Included antibody-coated microplate strips were removed and 50-100 µL of each standard, blank, and sample was added to the appropriate cells. Antibody was added to the wells for 1 hour (mouse) or 3 hours (human). Sample wells were washed with included wash buffer 3x. Specific for the human ELISA, a secondary streptavidin-HRP antibody was added to the wells for 30 minutes, and then washed as previously described. TMB solution was added to each well of both ELISAs for 10 minutes shaking at 400 rpm in the dark. Stop solution was added to each well and the absorbance at 450 nm was read immediately. The data was plotted and analyzed using GraphPad Prism and a one-way ANOVA.

### Cell Death Quantification by LDH Measurement

To quantify the viability of cells pre/post-treatment, the amount of lactate dehydrogenase (LDH) secreted into the media was measured. This was done using the CytoTox 96 Non-Radioactive Cytotoxicity Assay (Promega) per the manufacturer’s instructions as previously described^71^. Briefly, a 50 µL aliquot of all cell samples and a no-cell control were added to a 96-well plate and incubated with 50 µL CyTox 96 Reagent in the dark for 30 minutes at room temperature. After the incubation, 50 µL of Stop Solution was added to each well and the absorbance was read at 490 nm. The percentage of toxicity was calculated based on the Maximum LDH release controls and plotted and analyzed using GraphPad Prism and a one-way ANOVA.

### Co-Immunoprecipitation of NLRP3 inflammasome components

Immortalized human THP1 or mouse BMDMs were activated and treated with various small moelcules as described above. Cell were lysed gently using 300 µL ice cold 0.025M Tris pH 7.4, 0.15M NaCl, 0.001M EDTA, 1% NP40, 5% glycerol, and protease/phosphatase inhibitor and rotating at 4c for 5 minutes. The lysate was clarified by spinning at 13,000 x g for 10 minutes and the supernatant was saved for protein concentration determination using bradford assay (abcam ab102535) and co-immunoprecipitation. 100 µg of each sample was used for analysis, with the total volume normalized to 100 µL. And NLRP3 capture antibody (ABclonal A12694) was added to each sample at a 1:200 ratio. The samples were mixed at room temperature for 1 hour. While mixing, 50 µL of protein G beads (Millipore Sigma LSKMAGG10) per sample were washed three times with 500 µL PBS +0.01% Tween using a DynaMag-2 magnet. The beads were resuspended in PBS + 0.01% Tween such that an equal amount of bead suspension (100 µL) could be added to the antibody/sample mixture. This new mixture was rotated at room temperature for 1 hour. After incubating, samples were pulled back onto the magnet, the supernatant/unbound fraction was removed, and the beads were washed again three times with 500 µL PBS +0.01% Tween. Samples were eluited off the beads by adding 15 µL of 4x sample buffer (NuPage) + 6 µL of 10x Reducing agent (NuPAGE) + 39 µL DEPC water and heating for 10 minutes at 80C. The boiled samples were placed on magnet to pull back the beads the supernatant was loaded onto a NuPAGE™ 4 to 12%, Bis-Tris 1 mM 15-well mini-gels at 200 V for 30 minutes. Samples were transferred to PVDF membranes, blocked with 2.5% BSA in TBST, and probed with a primary antibody against NLRP3 (Adipogen), NEK7 (abcam), ASC (Adipogen), or pro-capspase-1 (CellSignaling). Blots were incubated with an HRP-linked secondary antibody (either mouse, rabbit, or goat depending on the species of the primary) and imaged using the iBright 1500 Imaging system. The intensities of the bands were quantified using the iBright Analysis Software. The intensity values were plotted and analyzed using GraphPad Prism and a one-way ANOVA.

### Immunofluorescence assays

Immortalized mouse BMDMs were plated into 12-well plates containing poly-l-lysine (NC9663893; Fisher Scientific) coated coverslips and left to adhere overnight. Cells were treated as previously described in the methods. After cell treatments, cells were washed with phosphate-buffered saline (PBS) to remove medium before cells were fixed with 4% paraformaldehyde (PFA) (sc-281692; Santa Cruz Technology) for 30 minutes. Coverslips were washed with PBS again and permeabilized with 0.2% Triton X-100 for 10 minutes while on ice. After blocking with 0.5% bovine serum albumin (BSA) blocking buffer for 30 minutes, coverslips were incubated with primary antibodies (diluted in 0.5% BSA blocking buffer) overnight at 4°C. Primary antibodies used were for NLRP3 (AdipoGen, Cryo2 AG-20B-0014-C100) and ASC (AdipoGen AL177). The following day, coverslips were washed before being incubated with secondary antibodies (1:5000, diluted in 0.5% BSA blocking buffer) for 40 minutes at room temperature. Finally, coverslips were mounted onto glass slides with ProLong Gold antifade reagent with 4′,6-diamidino-2-phenylindole (DAPI) (P36931; Invitrogen) to stain cell nuclei. Images were acquired on Zeiss LSM-900 microscope with Airyscan using 20× magnification. The excitation lasers used to capture the images were 488, 568, 630 and 405 nm using Alexa 488–, Alexa 568–, and Alexa 647–conjugated secondary antibodies. The same brightness/contrast profile was applied to all images within the same experiment. Five to ten images were captured per condition, where each image was considered a field. Zeiss and ImageJ imaging software were used for image analyses (an average of 10 cells per field, depending on magnification and cell type). Roughly 150-200 total cells were quantified per condition.

### Isolation and quantification of iBMDM and primary human PBMC RNA by reverse transcription PCR (RT-PCR) and Real-Time PCR (qPCR)

Immortalized iBMDM cells or primary human PBMC cells were treated in the presence of OGG1 small molecules as described above. Cells were washed with pre-chilled PBS and harvested at 1000 xg for 5 min. The total RNA was isolated from the cells using the RNeasy Mini RNA Purification kit (Qiagen) per the manufacturer’s instructions. The RNA concentration was checked via A260 absorbance and 5 µg of each sample was used in the reverse transcription reaction using the iScript Advanced cDNA Synthesis Kit (Bio-Rad) per the manufacturers instructions. Real-time PCR reactions were prepared with primers against *nlrp3*, and *asc*, along with *gapdh* as a control. Reactions were run using the SsoAdvanced Universal SYBR® Green Supermix (Bio-Rad). The qPCR reactions were run on the CFX Duet Real-Time PCR machine using the following conditions Initial denaturation at 98°C for 3 min, then 40 cycles of denaturation at 98°C for 15 s and annealing at 60°C for 30 s. This was followed by a melt curve from 65°C to 95°C with 0.5°C steps and 5 s per step. The readout was visualized using the BioRad CFX Maestro Software. The relative amounts of mRNA of either NLRP3 or ASC were calculated using GAPDH as a control. The final ΔΔCq were plotted and analyzed using GraphPad prism and a one-way ANOVA.

### Isolation of THP1 cytosolic and nuclear DNA and Real-time PCR (qPCR) analysis of nuclear DNA

Wild-type human THP1s were treated in the presence or absence of hOGG1 small molecules as described above. Briefly, cells were washed with pre-chilled PBS and harvested at 1000 xg for 5 min. The pellet was resuspended in a hypotonic lysis buffer(HLB) (10mM Tris HCl, pH 7.5, 10mM NaCl, 3mM MgCl2, 0.3% NP-40, 10% glycerol) and incubated rotating at 4 °C for 10 minutes. The mixture was then centrifuged for 2 minutes at 200 xg. The supernatant was saved as the cytosolic fraction in new tubes, the remaining nuclear pellet was resuspended in HLB and centrifuged again for 2 minutes at 200 xg to wash. This step was repeated two more times. The washed pellet was then resuspended in nuclear lysis buffer (NLB) (20mM Tris HCL, pH 7.5, 150mM KCL, 3mM MgCl2, 0.3% NP-40, 10% glycerol), vortexed, and lysed with a 18 gauge needle syringe by pulling up and expelling the liquid 10 times on ice. Both the tube with the pre-saved cytosolic and the newly lysed nuclear fraction were centrifuged for 15 minutes at 20,000 xg. The supernatant from each was collected into new tubes and used for subsequent DNA purification. The cytosolic and nuclear fractions of DNA were purified using the AllPrep DNA/RNA Mini Kit (Qiagen) per the manufacturer’s instructions. After purification was completed, the DNA concentration was evaluated by reading the A260 in a nanodrop. qPCR reactions were set-up with primers for *hTert* (nuclear DNA), *dDLoop* (mitochondrial DNA), and *GAPDH* (control) using the SsoAdvanced Universal SYBR Green Supermix (BioRad # 1725270) per the manufacturer’s instructions. The qPCR reactions were run on the CFX Duet Real-Time PCR machine using the following conditions: Initial denaturation at 98°C for 3 min, then 40 cycles of denaturation at 98°C for 15 s and annealing at 60°C for 30 s. This was followed by a melt curve from 65°C to 95°C with 0.5°C steps and 5 s per step. The readout was visualized using the BioRad CFX Maestro Software. The Cq values obtained for GAPDH DNA abundance served as normalization controls for the DNA values obtained from the test genes. The final ΔΔCq from the cytosolic and nuclear fractions of *hTert* were compared and plotted and analyzed using GraphPad prism and a one-way ANOVA.

### Purification of full-length NLRP3 decamer bound to TH5487

Wild-type NLRP3 was cloned into the mammalian expression vector pcDNA3.1HisB. The plasmid was expressed in DH5α cells (New England Biolabs) and purified using the PureLink HiPure Plasmid Maxiprep Kit (Thermo Fisher). The protein was expressed using the Expi293 Expression System (Thermo Fisher) in the presence of 0.01 mM TH5487 in DMSO. Briefly, cells were grown in Expi293 expression media until they reached a concentration of 3×10^6^ cells per milliliter and sustained viability of ≥95% live cells. At that time, 1 μg of expression vector was transfected per every 1 mL of cells with Expifectamine reagent. Once the cells reached viability of ≤80% live cells, they were harvested by spinning at 300 rpm for 5 min. The supernatant/dead cells were aspirated from the top, and the pellet was washed with cold PBS. Cells were resuspended in lysis buffer containing 50mM Tris, 150mM NaCl, 0.5mM TCEP, 10mM MgCl_2_, 0.01 mM TH5487, 1 mM ADP, 0.1 mM PMSF pH 7.5. The suspension was lysed by sonication with 5 seconds on, 10 seconds off, for a total of 4 minutes at 40%. The soluble lysate fraction was isolated by centrifuging at 100,000 xg for 1 hour and further clarified by filtering through at 0.45 um syringe filter. A HisTrap FF crude 5 mL column was equilibrated in binding buffer (50mM Tris, 150mM NaCl, 10mM MgCl_2_, 5% glycerol, 0.01mM Th5487, 1 mM ADP, pH 7.5). After the sample was loaded onto the column, it was washed with 10 column volumes (CV) of the wash buffer above, then eluted with 250 mM imidazole using a 50% gradient of the buffer 50mM Tris, 150mM NaCl, 10mM MgCl2, 5% glycerol, 500mM imidazole, 0.01mM Th5487, 1 mM ADP, pH 7.5. Peak fractions were pooled and the affinity purified protein was crosslinked using 0.5 mM bis(sulfosuccinimidyl)-suberate (BS3) for 30 minutes at 4 °C. The reaction was quenched by addition of 100 mM ammonium hydrogen carbonate for 15 min at 4 °C. The cross-linked protein was further purified by size exclusion using a HiLoad 16/600 Superose 6 pg size exclusion column equilibrated in binding buffer. Peak fractions were concentrated to ∼10mg/mL and visualized by SDS and Native gels, and western blots against NLRP3.

### Streptavidin-affinity grid preparation and NLRP3 complex visualization using cryoEM

NLRP3:TH5487 decemer was mixed with and boit-non-oxDNA and incubated on ice for 1 hour. The prepared grids are not glow-discharged. A total of 4 µL l of sample was added to the non-glow discharged grids containing a lipid-biotin layer. The grids were placed in the humidity chamber to make them hydrophilic for 5-10 minutes. The grids were then inverted and touched to wash buffer (50mM Tris, 150mM NaCl, 10mM MgCl_2_, 5% glycerol, 0.01mM Th5487, 1 mM ADP, pH 7.5) 1-3 times. The after touching the grid to buffer, the grid was then touched to 0.01% beta-octyl-glucopyranoside (BOG) in wash buffer was added. Then, the grid was released in a 20 µl 0.01% BOG drop, picked up with plunging tweezers, inserted into a Leica plunger, and blotted once only on the side with the sample. An additional 4 µL 0.01% BOG added to the top of the grid. The sample was blotted again, plunged, clipped and loaded onto a Tital Krios microscope equip with a Falcon 4i camera. A total of 26,390 movies were collected at 0.743 Å /pix at a total dose of 40 e/Å^2^, an exposure time of 3.25 seconds, and 37 frames per movie across a defocus range of −1.5 to 2.5 µm. After collection, the movies were motion corrected using RELION MotionCore2, and the streptavidin lattice was subtracted from all micrographs using MatLab as previously described. Corrected micrographs and FFT were visualized using EMAN2.

### Surface plasmon resonance evaluations of NLRP3 and ox-mtDNA

The ligand conjugate (biot-ox-mtDNA) was injected to on an active flow cell coat over the surface of an 8-channel Cytiva Biacore chip at a flow rate of 10 µL min^-1^. One channel in the flow cell was unmodified so serve as a control. The analyte (NLRP3) was diluted to 25, 20, 100, and 200 nM and injected at all 4 different concentrations at a flow rate of 30 µL min^-1^ for 180 seconds. The dissociation was then evaluated for 600 seconds. Experiments were run in triplicate. Data were collected using dual detection at 10 Hz and analyzed with the Biacore S200 Evaluation Software. The data was fit to both a 1-site binding model and a 2-side binding model, and the Chi^2^ values were used to define the best fit model.

### NLRP3 bound to TH10785 model generation using SWISS-MODEL

As previously described^12^, the amino acid sequence of hOGG1 that aligned with the NLRP3 pyrin domain (249-325) was saved as a ‘.pdb’ file from the published structure of hOGG1 bound of ox-DNA (PDBID: 1EBM). The sequence was uploaded to SWISS-MODEL as the User Template Modeling template file^72^. Then, various truncations of the NLRP3 pyrin domain were loaded as test target files to see which generated a viable model. Empirically, NLRP3 amino acids 1-85 were chosen. The SWISS-MODEL projection produced one model of NLRP3 based on the template hOGG1 structure. This model was further analyzed using ChimeraX to compare it to the published structures of wild-type NLRP3 (PDB1D: 7PZC) and hOGG1 bound to activator TH10785 (PDBID: 7AYY)^73,74^.

### Quantification and statistical analysis

A one-way ANOVA and non-linear regression were used to conduct all statistical analyses herein. All statistical analyses were performed as indicated in the figure legends, where N represents the number of replicates. Bar graph data were presented as mean ± SEM. Line graph data were presented as mean +/- SD. P-values <0.05 were considered statistically significant.

## Notes

### Competing Interest Statement

HMH has consulted for Novartis, Sobi, Ventyx, and Akros and has had research collaborations with Jecure, Zomagen, Takeda, and Inapill. The University of California Irvine is in the process of filing a patent with AL, JEC, and RM listed as inventors.

